# Phase Separation and a Hydrodynamic Instability Localize Proteins at Growing Microtubule Ends

**DOI:** 10.1101/2025.04.11.648094

**Authors:** Joël Schaer, Julie Miesch, Valeria Catapano, Ludovic Dumoulin, Joshua Tran, Marie-Claire Velluz, Charlotte Aumeier, Karsten Kruse

## Abstract

Regulating microtubule dynamics is essential for cellular function, and precise localization of regulatory proteins at microtubule ends is critical. End-binding protein EB3, a key regulator of microtubule growth, accumulates over an extended region at the growing end, forming a comet that gradually fragments into transient droplets along the microtubule shaft. Here, we combine *in vitro* reconstitution experiments with theoretical analysis to show that surface-mediated condensation of EB3 effectively localizes the protein at microtubule ends. Our results reveal that a Rayleigh–Plateau instability limits the condensate’s extent, producing a finite comet length and discrete droplets along the shaft. Remarkably, the comet size is independent of the GTP–cap size. This finding is supported by experiments on cells showing that modulation of microtubule growth velocity and hence GTP–cap size do not consistently alter EB3 comet length. Furthermore, our theory shows that rapid droplet evaporation requires a transition of EB3 to a non–phase-separating state. Overall, our work challenges the view that comet length directly reflects GTP–cap size and highlights a novel mechanism for regulating microtubule dynamics.

---

The microtubule cytoskeleton is essential for many cellular processes, including division, migration, and intracellular transport. Throughout these processes, the microtubule network undergoes continuous rearrangement, with microtubule dynamics playing a critical role in these changes. Consequently, regulation of these dynamics must be finely controlled. To achieve this, cells rely on a wide range of proteins. Among these regulators, end-binding proteins (EBs) can modulate microtubule dynamics directly, by interacting with microtubule growing ends, and indirectly, by recruiting further proteins to the microtubule plus end [1–3].

Microtubules grow through the addition of guanosine triphosphate (GTP)-bound *αβ*-tubulin heterodimers. Within a microtubule, the GTP bound to *β*-tubulin hydrolyses to guanosine diphosphate (GDP), which is associated with a conformational change of the tubulin dimer [4, 5]. As a result, microtubules consist of GDP-tubulin along their shaft and exhibit a region of GTP-tubulin at the growing end. The size of this GTP-cap is determined by the ratio between the microtubule growth speed and the GTP hydrolysis rate. Presumably by reading out the tubulin conformation within the microtubule lattice, EBs assemble into a comet at the growing end (Fig. 1A, B, Movies S1, S2) [1, 6, 7]. For this reason, EB comets are routinely used *in vitro* and in cells as a proxy for the GTP-cap size [1, 7–9]. In agreement with this idea, the EB-comet size increases linearly with the microtubule growth speed *in vitro* [10]. In cells, reducing microtubule growth speed by application of low concentrations of the tubulin-targeting drug nocodazole reduces comet size [11]. However, also longer comets have been observed in association with slower-growing microtubules [12]. Furthermore and at odds with common models of GTP-hydrolysis [13], in cells, transient structures called remnants often detach from the comet, a phenomenon not seen *in vitro* (Fig. 1A, B, Movies S1, S2) [14–19]. Consequently, the link between EB comets and GTP caps is more complex than previously assumed.

**FIG. 1.**
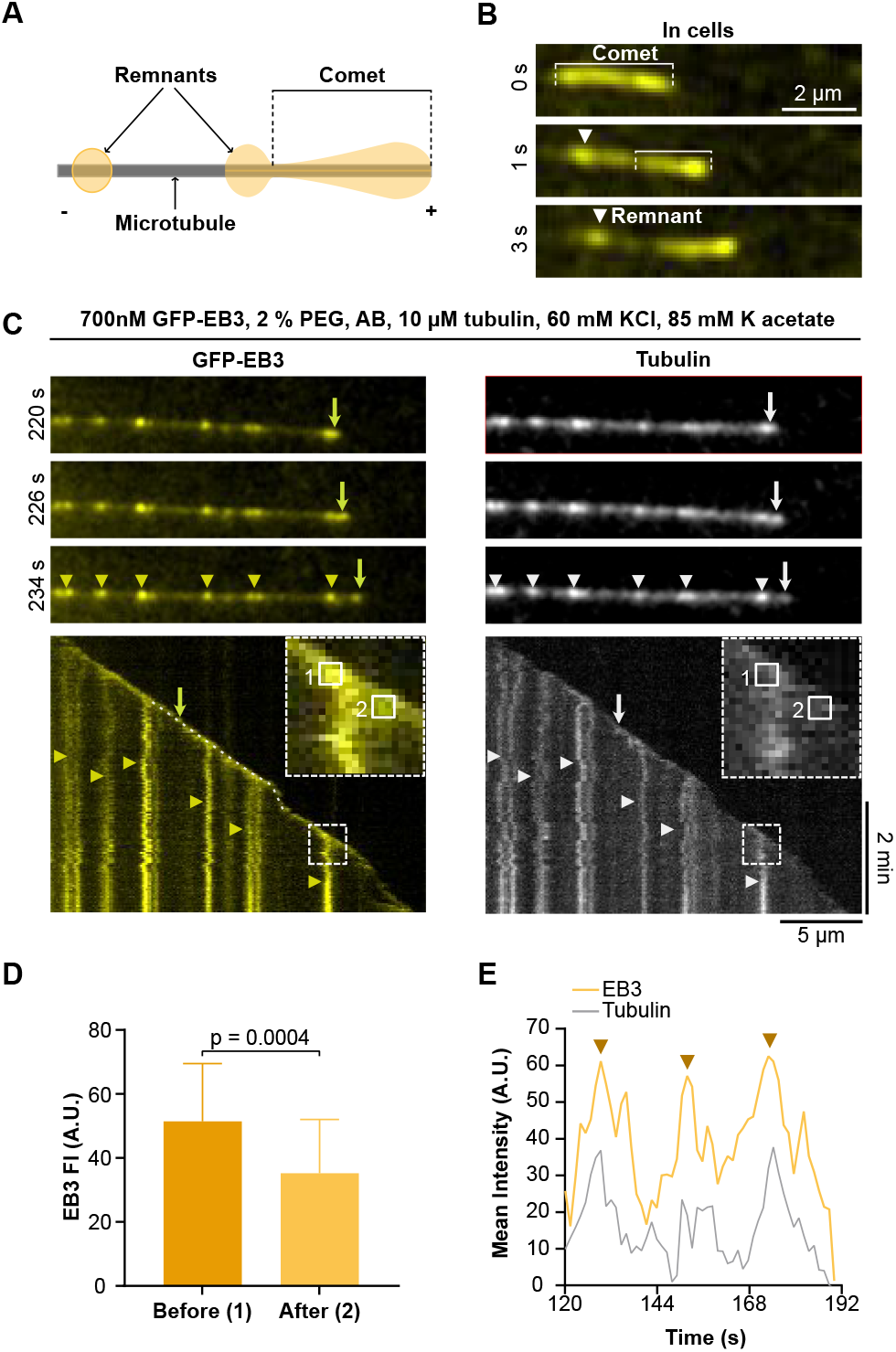
EB3 comets with remnants on growing microtubules. (A) Scheme of EB3 comets forming remnants behind the growing microtubule end. (B) Representative confocal time-lapses of C2C12 cells transfected with EB3-GFP showing the fission of a remnant from the comet (white arrowhead). (C) Top: representative TIRF snapshots of a growing microtubule *in vitro*; GFP-EB3 (left) and tubulin (right). Bottom: corresponding kymographs. Arrows indicate the EB3 comet, arrowheads remnants. Dashed rectangles indicate zoom-ins. Squares 1 and 2 indicate times when fluorescence intensities for panel (D) were taken. The dotted line indicates the linescan presented in (E). (D) GFP fluorescence of EB3 comets at the growing microtubule end just before and after remnant fission. Mean with SD from two independent experiments with 22 comets analyzed. (E) Fluorescence intensity at the growing microtubule end after subtraction of the background from the kymograph in (C) showing accumulation of tubulin and GFP-EB3 at the microtubule end. The arrowheads represent the fission events.

Given the unresolved relationship between EB comet length and microtubule growth dynamics, recent findings that EB proteins can undergo liquid–liquid phase separation (LLPS) [20–23] suggest an alternative mechanism that might reconcile these observations. Through LLPS, macromolecules form a dense phase coexisting with a dilute phase [24] and surfaces can act as nucleators during this process [25]. For example, stabilized microtubules provide a surface that promotes phase separation of the microtubule-binding protein TPX2, which forms a growing film that collapses into dense droplets regularly spaced along the microtubule [26]. Whether phase separation can contribute to the localization of EBs at growing microtubule ends remains unexplored. In the present study, we address this gap by combining experimental reconstitution with theoretical analysis.

We use a combination of *in vitro* experiments and theory to study the condensation of EB3 on growing microtubules. We find that liquid-liquid phase separation in combination with an instability of the interface between the dilute and the dense phases is sufficient for comet formation. Our theory shows that, in this scenario, the size of the GTP cap has only a weak effect on comet length. Experiments we performed in cells in which we modified the microtubule growth velocity are in line with this finding.

## *IN VITRO* RECONSTITUTION OF EB3 COMETS WITH REMNANTS

*In vitro* under non-phase separating conditions, EB3 interacts with the growing microtubule end by forming a comet-like structure without remnants [8, 27]. To elucidate the mechanism underlying both comet and remnant formation, we established an *in vitro* reconstitution system under phase-separating conditions. In our experiments, microtubules were grown from stabilized seeds in the presence of 10 *μ*M tubulin, 700 nM GFP-EB3, salt to reduce EB3 binding to the GDP-shaft (60 mM KCl, 85 mM KAc), and an anti-bleaching buffer (AB) to enable imaging for up to 45 min. The anti-bleaching buffer strongly reduced phase separation, increasing the EB3 threshold concentration for droplet formation (SI Appendix, Fig. S1A,B). To promote phase separation in this condition and to mimic the crowded cellular environment we added 2 % polyethylene glycol (PEG). We used total internal reflection fluorescence (TIRF) microscopy to visualize microtubule growth and the GFP-EB3 distribution.

Under these conditions, EB3 reliably tracked growing microtubule ends and formed comets. Behind the comets, distinct EB3 patches formed (Fig. 1C). They separated from the comet and then remained immobile on the microtubule while growing as indicated by their increasing fluorescence. Unlike cellular remnants that disassemble within seconds [22], the patches persisted for tens of minutes. No bulk phase separation of EB3 was observed during the experiments (SI Appendix, Fig. S1C).

In agreement with previous reports [22], tubulin was enriched within EB3 comets. Consistently, tubulin was also enriched within EB3 remnants along the microtubule shaft (Fig. 1C). Notably, EB3 fluorescence intensity increased over time until the comet reached a threshold intensity, which correlated with triggering remnant fission (Fig. 1D). Similarly, tubulin intensity increased at the microtubule end and then decreased with remnant fission (SI Appendix, Fig. S1D). Interestingly, tubulin accumulation at the end happened only after an initial increase of EB3 fluorescence (Fig. 1E). Together, these observations confirm that our *in vitro* system can faithfully reproduce EB3 comet formation with distinct remnants under phase separating conditions.

## PHYSICAL DESCRIPTION OF A PHASE-SEPARATING MIXTURE ON A CYLINDRICAL SURFACE

To quantitatively capture the physical principles underlying EB3 comet and remnant formation, we developed a continuum description of phase separation in presence of a cylindrical surface. In this description, EB3 is the solute and the buffer is the solvent, while the microtubule is represented by a cylinder. This framework provides a means to connect the binding properties of EB3 to the macroscopic features observed in our experiments.

### Phase-field Description of a Binary Mixture

Consider a solute like EB3 that phase separates into a dense phase (concentration *c*_den_) and a dilute phase (concentration *c*_dil_). Let *c*(**r**) denote the solute concentration at position **r**. We define the phase field *ϕ* as

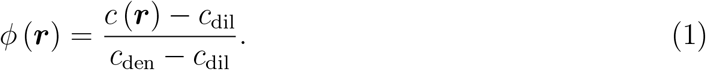

which is the local excess concentration relative to the dilute phase normalized by the excess concentration of the dense phase. The phase-separation dynamics is given by the Cahn-Hilliard equation

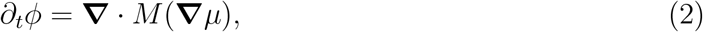

where *M* is the solute mobility and *μ* = *δ*Ψ/*δϕ* is the exchange potential, where the free energy Ψ is given by

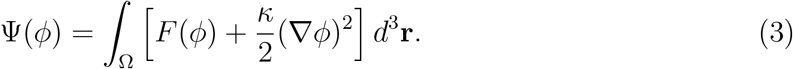

Here, Ω is the volume of the system and the free energy *F* (*ϕ*) = *βϕ*^2^(*ϕ* − 1)^2^, which is akin to the Flory-Huggins free energy [28, 29], but is numerically more stable. In both cases, the potential has two minima, such that the dynamics is qualitatively similar. The exchange potential is given by

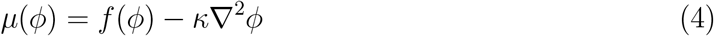

with 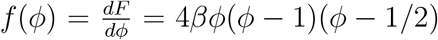.Further, we relate the abstract parameters *β* and *κ*, to the physical parameters interfacial tension *σ* and capillary width of the interface *ϵ* by 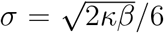 and 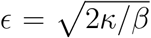 [30, 31]. Note that the parameters *M* and *σ* only appear in form of the product *Mσ* in the Cahn-Hillard equation 2.

The dynamic equations 1-4 describe phase separation of the binary mixture for *β* > 0 if the average phase-field *ϕ*_0_ lies between 0 and 1, that is, *c*_dil_ < *c*_0_ < *c*_den_, where *c*_0_ is the mean concentration. At equilibrium, the minority phase forms a single spherical droplet within the majority phase. In summary, this phase-field description provides a mathematical framework to describe the kinetics and thermodynamics of EB3 phase separation, setting the stage for our investigation of surface effects in this context.

### Interaction of the Phase Field with a Surface

In order to study in our phase-field description how binding of EB3 to the microtubule surface protein affects phase-separation dynamics, we introduce appropriate boundary conditions. Concretely, we describe a microtubule as a solid cylinder with the radius of a microtubule, *R*_*i*_ = 12.5 nm. The interaction between the solute and the cylinder surface *S* is captured by a surface energy

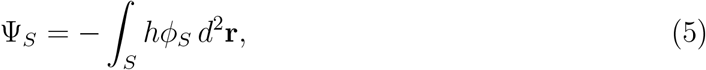

where *ϕ*_*S*_ is the phase field at the surface and *h* represents the interaction strength – positive for attraction and negative for repulsion. Depending on our questions the parameter *h* was varied along the surface to capture differences in the binding of EB3 to the GTP-cap and the GDP–shaft. Minimizing the total free energy, the contribution Eq. 5 yields the boundary condition

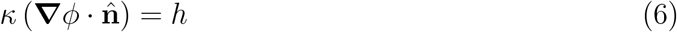

at the cylinder surface, where 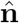 is its outward normal of *S*. We also impose no-flux boundary conditions at all surfaces, 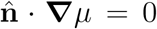.Thus, by incorporating surface interactions into our description, we relate the binding of EB3 on the microtubule to the resulting phase-separation dynamics. This completes the physical description of protein phase-separation in presence of a cylindrical substrate.

### Condensation at the Growing End Does Not Explain Comet Formation

We now apply our theoretical framework to the dynamic environment at the growing microtubule end. To capture conditions at the growing end, we solved the dynamic equation 2 numerically in an axisymmetric geometry (SI Appendix, Materials and Methods). The binary mixture is confined within a cylindrical reaction chamber of radius *R*_*o*_ = 500 nm, co-axial with the microtubule-representing cylinder. To mimic the GTP-cap of a microtubule, we set an attractive interaction (*h* > 0) along the cap-length ℓ_cap_ that transitions sharply to *h* = 0 for the GDP-shaft.

Depending on parameter values, our simulations produced two distinct solutions: i) A continuous film with a gradually increasing height *ξ* from the growing end (Fig. 2A, Movie S3) or ii) a periodic formation of droplets at the growing end (Fig. 2B, Movie S4).

**FIG. 2.**
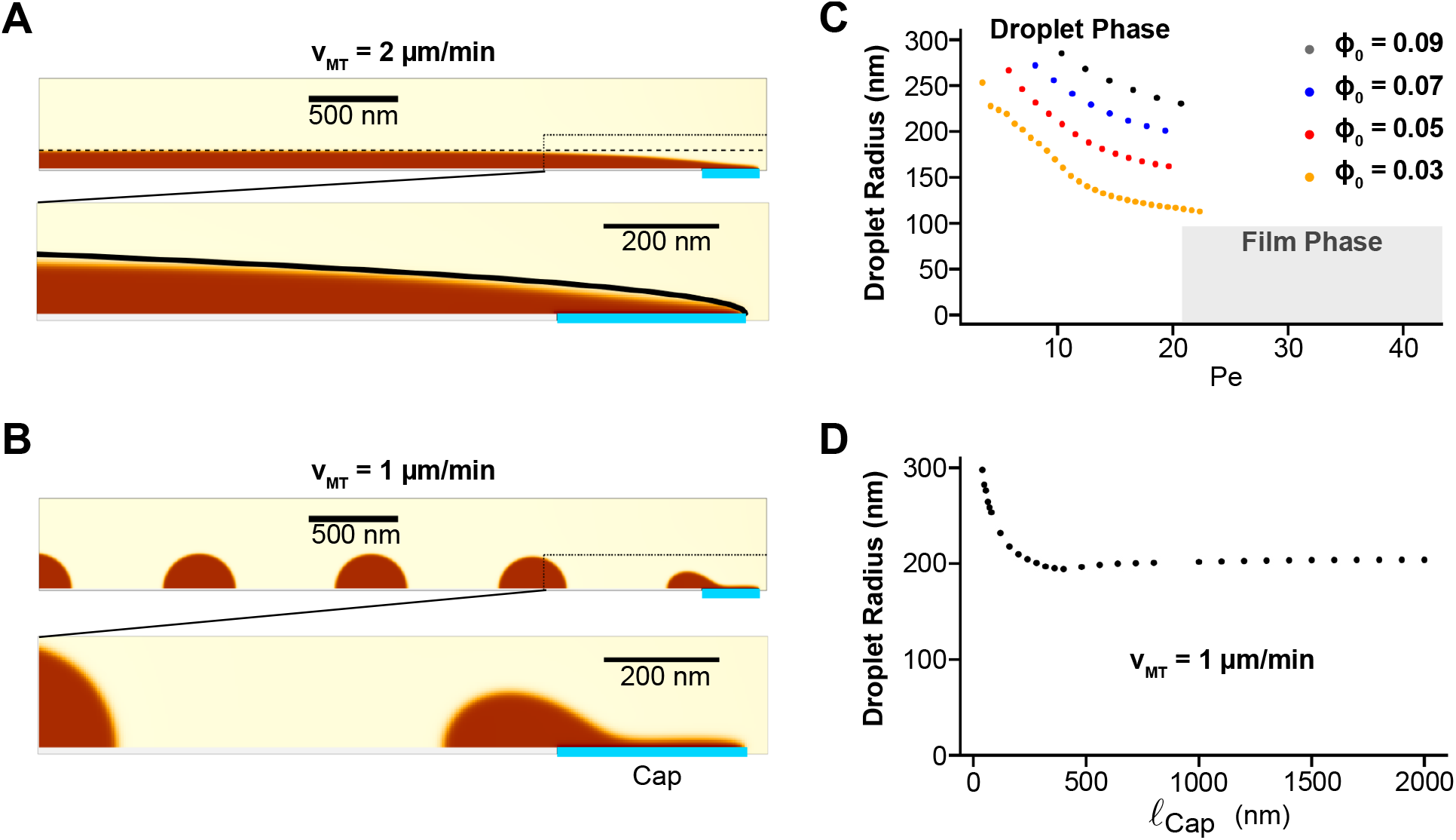
Physics of condensation on a growing cylinder. (A) Top: Steady-state phase field on a growing cylinder for Pe ≳20. Dashed line: saturated height *ξ*_∞_, Eq. 9. Dotted box: zoom-in region. Bottom: zoom-in on the microtubule end. Solid line: film height *ξ*, Eq. 7. (B) Top: Snap-shot of a phase field on a growing cylinder for Pe ≲ 20. Dotted box: zoom-in region. Bottom: zoom-in on the microtubule end. Cylinder growth velocity *v*_MT_ = 2 *μ*m/min (A) and *v*_MT_ = 1 *μ*m/min (B). Other parameter values as in (SI Appendix, Materials and Methods). (C) Droplet radius as a function of Pe when changing the cylinder growth velocity *v*_MT_ and for different bulk concentrations *ϕ*_0_. (D) Droplet radius as a function of ℓ_cap_. Parameter values as in SI Appendix, Materials and Methods.

For the continuous film solution (i), we computed the profile by analyzing the solute dynamics in a cross-section perpendicular to the cylinder axis. Assuming an infinite reaction chamber *R*_*o*_ = ∞ and setting *ϕ* = *ϕ*_0_ as |**r**| → ∞, we calculated the film height *ξ* measured from the center of the cylinder to the interface between the dense and the dilute phases over time [26] (SI Appendix, Computation of the film profile). For a constant growth velocity *v*_MT_, we find

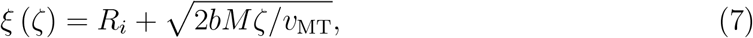

where *ζ* is the distance along the cylinder surface from the cylinder end. The dimensionless constant *b* is determined by

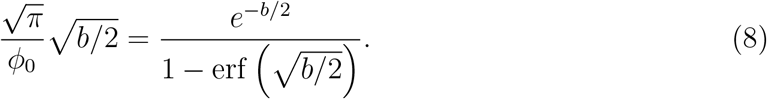

Our analytical result matches the interface profile of our numerical solution close to the cylinder end (Fig. 2A).

For a finite reaction chamber, *R*_*o*_ < ∞, the film height eventually saturates and reaches an asymptotic value which is determined by the total amount of solute

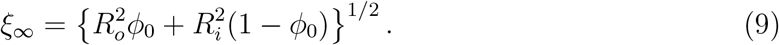

The results of our numerical solution agree well with this value (Fig. 2A).

For the droplet solution (ii), the droplets eventually form a contact angle of *π*/2 with the cylinder surface due to *h* = 0 along the shaft. Changing the interfacial tension *σ* or mobility *M*, while keeping *Mσ* = *const*, has no effect on the average droplet radius (SI Appendix, Fig. S2A). This is consistent with Eq. 2 that depends only on *Mσ*. Increasing *Mσ* leads to larger droplets due to faster diffusion (SI Appendix, Fig. S2A,B). In contrast, increasing the cylinder growth velocity *v*_MT_ yields smaller droplets, because faster growth promotes faster separation of a droplet from the end and thus faster nucleation of a new droplet. The newly nucleated droplet competes with the one formed previously for the available material (SI Appendix, Fig. S2C). In contrast, increasing the average phase-field value *ϕ*_0_ leads to larger droplets, because more material is available (SI Appendix, Fig. S2D).

Concerning the influence of the cap length ℓ_cap_ on the surface condensate, the droplet radius decreases for increasing ℓ_cap_ until ℓ_cap_ is approximately equal to twice the droplet radius, ℓ_cap_ ∼ 400 nm (Fig. 2D). For longer caps, the droplet radius remains constant. For continuous films, the asymptotic height *ξ*_∞_ is always independent of ℓ_cap_ (Eq. 9). In addition, we characterize the transition between films and droplets by forming a dimensionless number

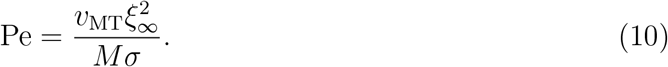

It is reminiscent of the Péclet number, which compares convection to diffusion. Our numerical results show that the film solution is stable for Pe ≳ 20, whereas droplets form for lower Pe values (Fig. 2C). Although Pe accounts for microtubule properties such as radius and growth velocity, it is independent of the nucleotide state along the microtubule, that is, ℓ_cap_.

In summary, our theoretical analysis demonstrates that the interplay between phase separation and surface interactions on a growing cylindrical substrate can yield condensation at the growing end in two distinct ways, either as a continuous film or as periodic droplets. However, condensation alone cannot explain the formation of EB3 comets with remnants at the growing end (Fig. 1C). Furthermore, our analysis suggests an unexpected independence between comet length and cap size [8].

### EB3 Droplets on Growing Microtubules without Comet Formation

We next sought to determine whether the different condensation patterns obtained in our theory could be recapitulated experimentally by altering EB3’s binding to the microtubule lattice. In our *in vitro* system, the length of the GTP-cap is intrinsically linked to the microtubule growth speed and the GTP-hydrolysis rate. Although we cannot tune the hydrolysis rate independently of the growth speed, we can override the cap’s attractiveness by modulating the salt concentration. Specifically, by reducing salt conditions to 60 mM KCl and no KAc, we enhanced the overall attractive interaction of EB3 with the microtubule. We also omitted PEG and increased the EB3 concentration. At 10 *μ*M EB3 (10% GFP-EB3, 90% EB3) and in the presence of 10 *μ*M tubulin, EB3 decorated the growing microtubule shaft with bright, stationary patches (Fig. 3A). Over time, the patches became brighter, indicating that EB3 continued to accumulate and, with it, tubulin. FRAP measurements confirmed that these patches were liquid droplets (Fig. 3B).

**FIG. 3.**
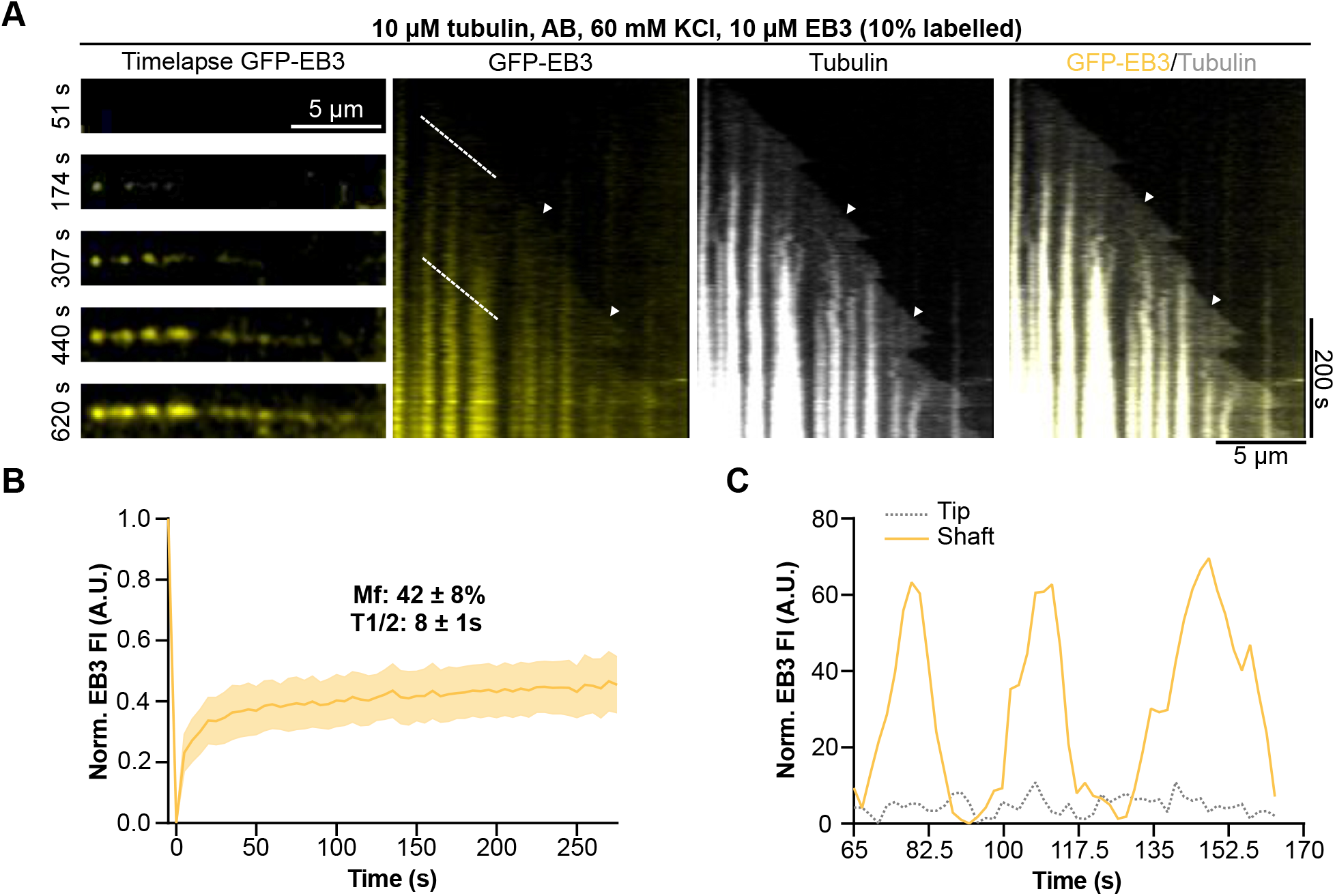
EB3 droplet formation on growing microtubules *in vitro*. (A) Representative TIRF images of GFP-EB3 (left) and corresponding kymographs of GFP-EB3 alone, tubulin alone, and the merge (right). Arrowheads highlight the end of the microtubule. (B) FRAP recovery curve of GFP-EB3 condensates on dynamic microtubules. The yellow curve shows the mean with SD of three individual experiments with a total of 39 condensates analyzed. (C) Scans along white lines of the kymograph in (A) corresponding to positions at fixed distances from the growing microtubule end showing strong accumulation of GFP-EB3 on the microtubule shaft over time, but only faint signal at the growing end.

Compared to the EB3 droplets on the microtubule shaft, the fluorescence of GFP-EB3 at the growing end remained relatively dim (Fig. 3C). No comets were observed in this condition. This behavior is consistent with our computations showing periodic droplet formation at the growing end when Pe ≲ 20 (Fig. 2B). However, for low salt concentrations, we did not observe a continuous film. While this appears to be inconsistent with our theory (Fig. 2A), previous work has shown that protein films growing on microtubules are prone to the Rayleigh-Plateau instability [26]. In this scenario, a growing film becomes unstable and breaks into discrete droplets, which would reconcile our experimental and theoretical results.

## THE RAYLEIGH-PLATEAU INSTABILITY LEADS TO COMET FORMATION

To determine whether the Rayleigh-Plateau instability contributes to droplet formation by fragmenting a continuous EB3 film, we extended our theoretical description to include fluid dynamics driven by interfacial-tension gradients.

### Fluid Dynamics of the Binary Mixture

To study flows in the binary mixture, we introduce the center-of-mass velocity field **u**. We assume the mixture to be incompressible such that

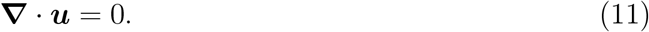

The velocity field **u** is determined by the Navier-Stokes equation

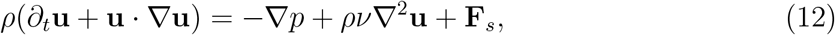

where *ρ* is the fluid mass density, *p* the hydrostatic pressure, *ν* the shear viscosity, and **F**_*s*_ a force that accounts for interfacial tension effects,

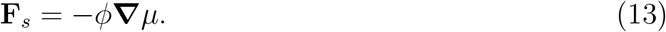

This force is obtained by taking the derivative of the interfacial stress 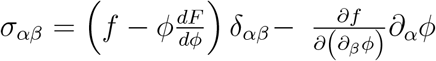 [32]. We solve Eq. 12 numerically using a Lattice-Boltzmann method (SI Appendix, Materials and Methods). Given that fluid flows are slow, the corresponding Reynolds number Re is small. We used parameters such that Re ≲1.

In the presence of a fluid flow, an advection term needs to be added to the Cahn-Hilliard equation 2,

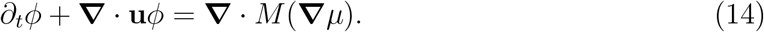

We imposed no–slip boundary conditions, **u** = 0, at all surfaces.

### Droplet Formation on Static Cylindrical Surfaces

We numerically studied the Rayleigh-Plateau instability of a condensing film first on a static cylindrical surface, with periodic boundary conditions along the cylinder axis. Consistent with Ref. [26], our description produced evenly spaced droplets along the cylinder surface (Fig. 4A, Movie S5).

**FIG. 4.**
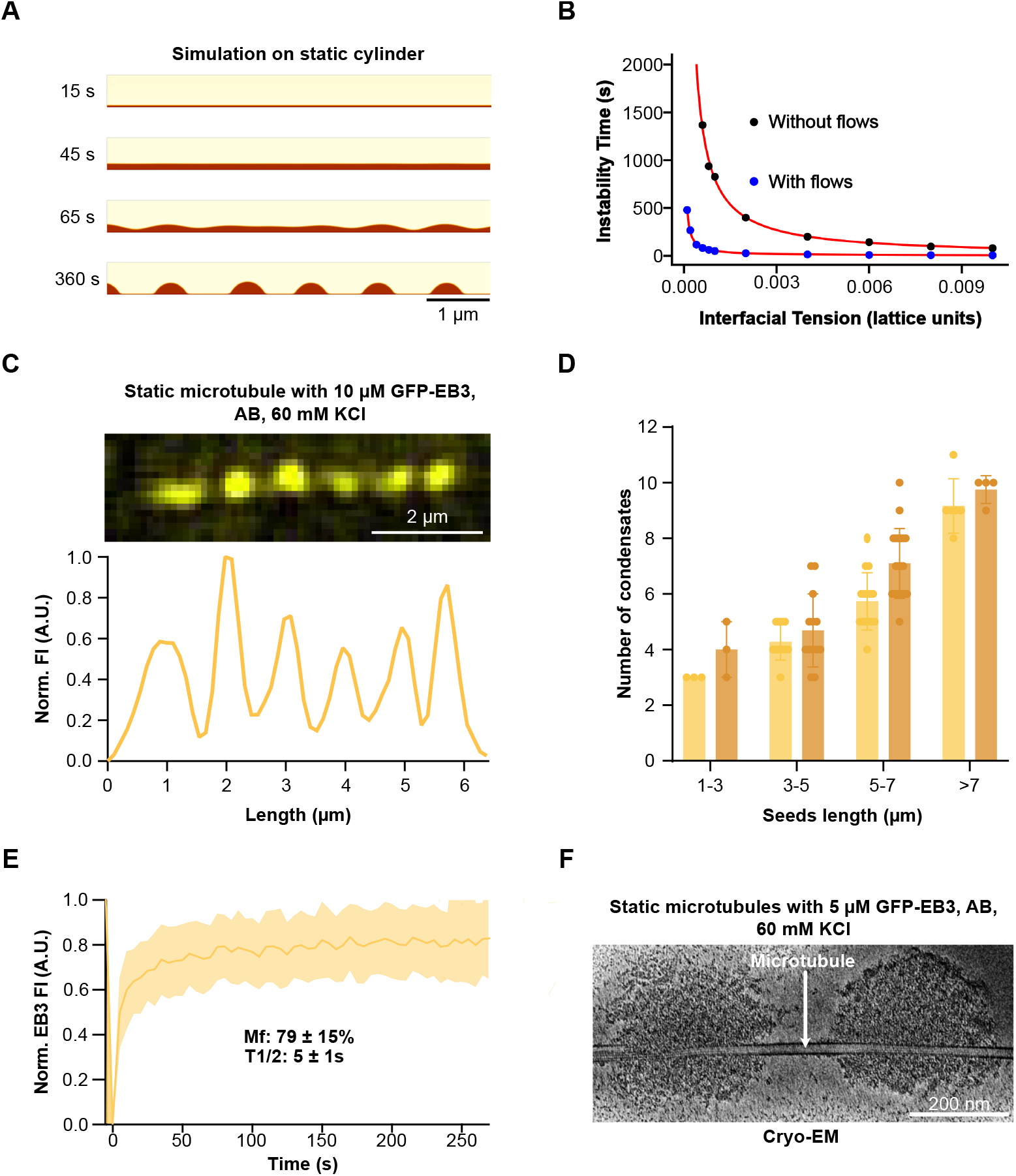
Droplet formation on static microtubules. (A) Snap-shots from a simulation showing condensation on a static cylinder. (B) The scale *τ*_*m*_ as a function of the surface tension *σ* with (blue) and without (black) hydrodynamic flows. Red lines are guides to the eye of the form ∼ 1/*σ*. Parameter values as in SI Appendix, Materials and Methods. (C) Top: Representative TIRF image of GFP-EB3 decorating GMPCPP-taxol microtubules. Bottom: Corresponding linescan of the fluorescence intensity over microtubule length. (D) Number of EB3 droplets for different seed lengths and 10 *μ*M (beige) and 20 *μ*M (brown) GFP-EB3. (E) FRAP recovery curve for GFP-EB3 droplets on a static microtubule. (F) Representative cryo-EM image of EB3 droplets on GMPCPP-taxol microtubules at 5 *μ*M GFP-EB3.

Two time scales are relevant in this context: the scale *τ*_*f*_ is determined by the growth rate of the film on the surface, whereas the scale *τ*_*m*_ is associated with the growth of the most unstable mode driven by the Rayleigh-Plateau instability. We determined these timescales by tracking the interface height along the cylinder axis. Specifically, *τ*_*f*_ is defined as the time at which the interface reaches 95% of its final height. This time is primarily governed by the mobility *M* (SI Appendix, Fig. S3A,B). In contrast, *τ*_*m*_, which is the time when the standard deviation of the interface height reaches 5% of the initial height, is largely determined by the interfacial tension *σ* (Fig. 4B) and viscosity *ν* (SI Appendix, Fig. S3C,D). For the parameters in Fig. 4A, the film grows faster than the unstable mode, *τ*_*f*_ < *τ*_*m*_.

We find *τ*_*m*_ ∼ 1/*σ* (Fig. 4B) and *τ*_*m*_ ∼ *ν* (SI Appendix, Fig. S3D) if *τ*_*f*_ < *τ*_*m*_, which is compatible with the analysis in Ref. [26]. Note that droplets form also in the absence of flows induced by interfacial-tension gradients. However, the latter can accelerate film breakup by more than an order of magnitude (Fig. 4B). These theoretical findings set the stage for our subsequent experiments on static microtubules, where we sought to confirm droplet formation driven by the Rayleigh–Plateau instability.

### EB3 Forms Droplets on Static Microtubules

To test the role of the Rayleigh-Plateau instability for EB3 experimentally, we studied EB3 behavior on double-stabilized microtubules (GMPCPP and taxol). At 10 *μ*M EB3, droplets formed along the entire microtubule (Fig. 4C), with fluorescence maxima separated by an average distance of 750±220 nm (SI Appendix, Fig. S4A). On longer microtubules, the number of droplets increased while the spacing remained constant (Fig. 4D) supporting the fact that they result from a finite-wavelength instability. Doubling the EB3 concentration increased the mean fluorescence intensity by more than two-fold (SI Appendix, Fig. S4B). In addition, the droplet spacing remained constant (SI Appendix, Fig. S4A). This is compatible with the results in Ref. [26], where the spacing between droplets varied by less than 50 nm as the film thickness doubled, as such small changes are inaccessible with the TIRF microscopy we use.

To probe the material properties of these fluorescent patches, we recorded fluorescence recovery after photobleaching (FRAP). Bleached patches recovered fluorescence with a half-time of 5 s and an immobile fraction of 21% (Fig. 4E). Furthermore, cryo-electron microscopy images showed that EB3 accumulated in spherical condensates around the microtubules (Fig. 4F). The spheres had radii ranging from 100 to 200 nm, with a thin EB3 layer on the microtubule between the spheres (Fig. 4F). This data validates our assumption of an axisymmetric EB3 distribution used in our numerical computations.

Together with our previous results on EB3 phase-separation [22] and with the spontaneous formation of EB3 condensates in solution at 10 *μ*M EB3 (SI Appendix, Fig. S4C), these results show that EB3 forms liquid droplets on static microtubules. The behavior closely resembles that of TPX2 [26] and supports our simulation results, which show that the Rayleigh–Plateau instability leads to fragmentation of EB3 films into droplets.

### Comets Form on Growing Cylindrical Surfaces in the Presence of Hydrodynamic Flows

Solving the dynamic equations 11-14 numerically for growing cylinders with boundary condition 6 again led to two types of solutions: i) periodic droplet formation at the growing end, and ii) a film extending from the growing end. However, whereas the film extended along the whole cylinder in absence of hydrodynamic effects, we now find that the film’s extension is limited by the Rayleigh-Plateau instability (Fig. 5A, Movie S6). Beyond the film, the cylinder is decorated with droplets, which separated from the film. This solution resembles the comets observed experimentally.

**FIG. 5.**
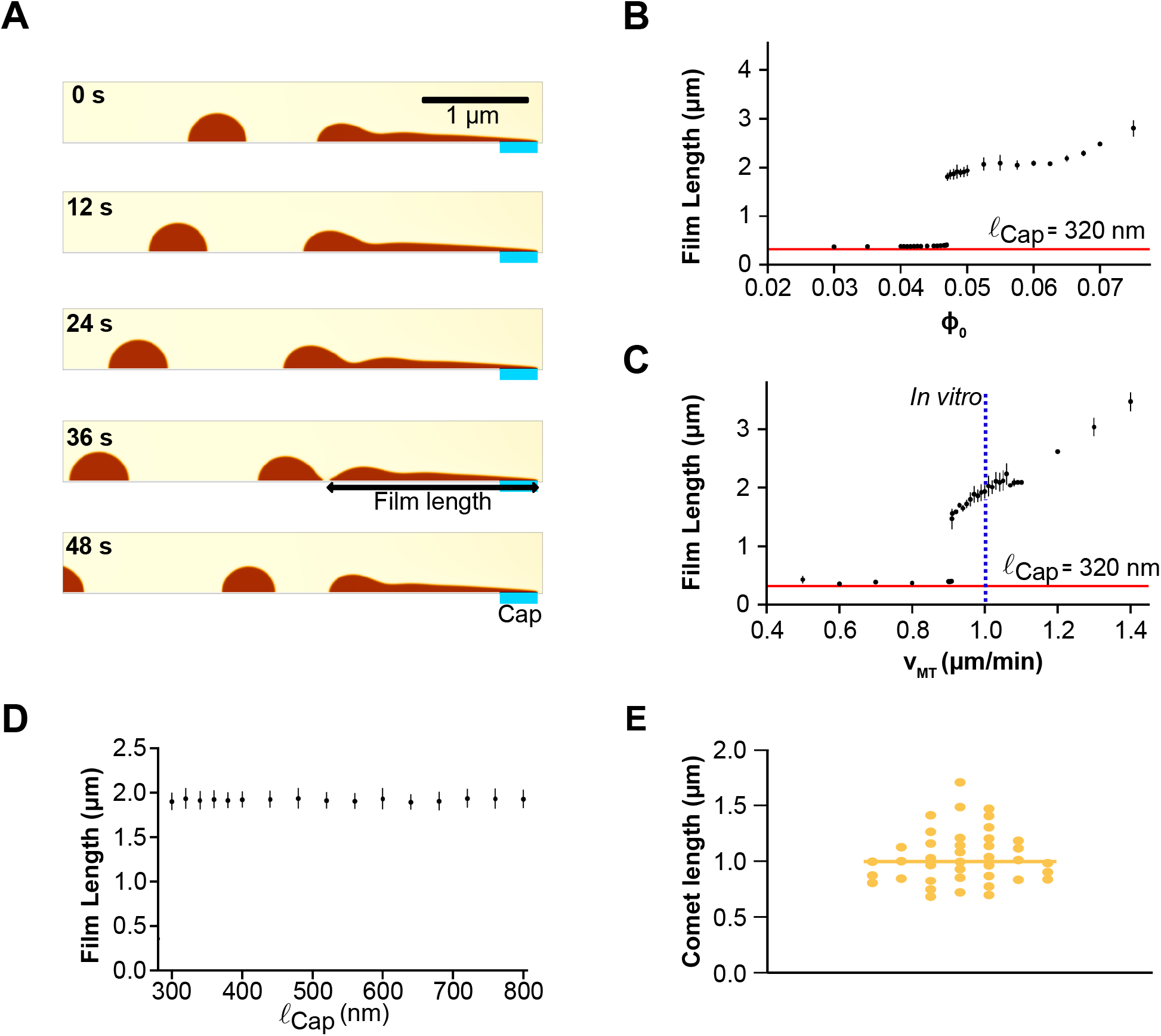
Comet formation at growing cylinder ends in the presence of hydrodynamics. (A) Representative snapshots from a simulation showing condensation on a growing cylinder with flows induced by interfacial gradients for ℓ_cap_ = 320 nm. (B-D) Film length as a function of the average bulk concentration *ϕ*_0_ (B), the cylinder growth velocity *v*_MT_ (C), and the cap length ℓ_cap_ (D). Red lines represent cap length ℓ_cap_ = 320 nm (B,C). Blue dotted line represents the average microtubule growth velocity observed *in vitro* (C). (E) Distribution of comet length *in vitro* under phase separating conditions (Fig. 1C) from 22 comets analyzed in two experiments. Parameter values as in SI Appendix, Materials and Methods except for *M* = 6.25 (A-D).

Increasing the bulk concentration *ϕ*_0_ leads to a longer film (Fig. 5B), because higher *ϕ*_0_ values accelerate film thickening, which in turn increases the timescale *τ*_*m*_ associated with the Rayleigh-Plateau instability [26]. Similarly, the film length increases with both, the cylinder growth velocity *v*_MT_ and the viscosity *ν* (Fig. 5C, SI Appendix, Fig. S5A). Notably, the film length increases with *v*_MT_ even when the cap length ℓ_cap_ is held constant (Fig. 5C), suggesting that the experimentally observed dependence of comet size on microtubule growth velocity is not solely due to variations in ℓ_cap_ as commonly assumed. At critical values of *ϕ*_0_ and *v*_MT_, however, the film length drops to ℓ_cap_ for smaller values of these parameters (Fig. 5B,C). Similar discontinuities appear at critical values of the viscosity, interfacial tension, and mobility (SI Appendix, Fig. S5). Contrary to the dependence on *ϕ*_0_ and *v*_MT_, the film length decreases with the interfacial tension *σ* and the mobility *M* (SI Appendix, Fig. S5B,C). Overall, our combined numerical and experimental data indicate that the Rayleigh–Plateau instability, when coupled with hydrodynamic flows, is a key factor for generating the EB3 comet structure observed *in vivo* and that the comet length decouples from the GTP-cap length at critical parameter values.

### The Comet Length is Independent of the GTP-cap Length

Our physical approach does not only allow us to recapitulate the organization of proteins at the growing microtubule end but also lets us address the longstanding assumption that comet length is directly linked to the length of the GTP-cap. While it is experimentally currently not possible to measure the cap length directly and to uncouple the growth speed reliably from the cap length, both are possible in our theory. It reveals that although nucleating a film requires a cap with *h* > 0, the resulting film length is independent of the cap length ℓ_cap_ for comet lengths in range with our *in vitro* experiments(Fig. 5D, E). In fact, even if *h* is uniformly positive along the entire cylinder, mimicking a microtubule in an all-GTP state, a film of finite length can still form (SI Appendix, Fig. S5D). This suggests that the length of an EB3 comet is not a good proxy for the GTP-cap length.

To test the connection between the comet length and the GTP-cap in a physiological environment, we manipulated the GTP-cap size in cells by reducing microtubule growth velocity with 50 nM nocodazole [11]. Nocodazole binds to free tubulin, reducing thereby the tubulin concentration available for polymerization. We grouped EB3-overexpressing C2C12 cells into low and medium EB3-GFP overexpression levels. In cells with low EB3-GFP levels, nocodazole halved the microtubule growth velocity from 8 *μ*m/min to 4 *μ*m/min (Fig. 6A,B). Since nocodazole interacts with free tubulin, GTP hydrolysis in microtubules is presumably unaffected by the drug and the GTP-cap length should decrease by the same factor. Compatible with this expectation, the comet length decreased from 2.2 *μ*m to 1.5 *μ*m. However, in nocodazole treated cells with medium-EB3-GFP levels, the mean comet length increased to 2.5 *μ*m, even though the microtubule growth velocity remained comparable to that of low-overexpressing cells (Fig. 6A,B). These results are consistent with our theoretical findings: changes in the GTP-cap size do not directly control EB3 comet length (Fig. 5D). Let us also note that for high overexpression levels, microtubules are completely covered by EB3-GFP (SI Appendix, Fig S6). This is consistent with the increase of the Péclet number Pe with *ϕ*_0_, Eq. 10.

**FIG. 6.**
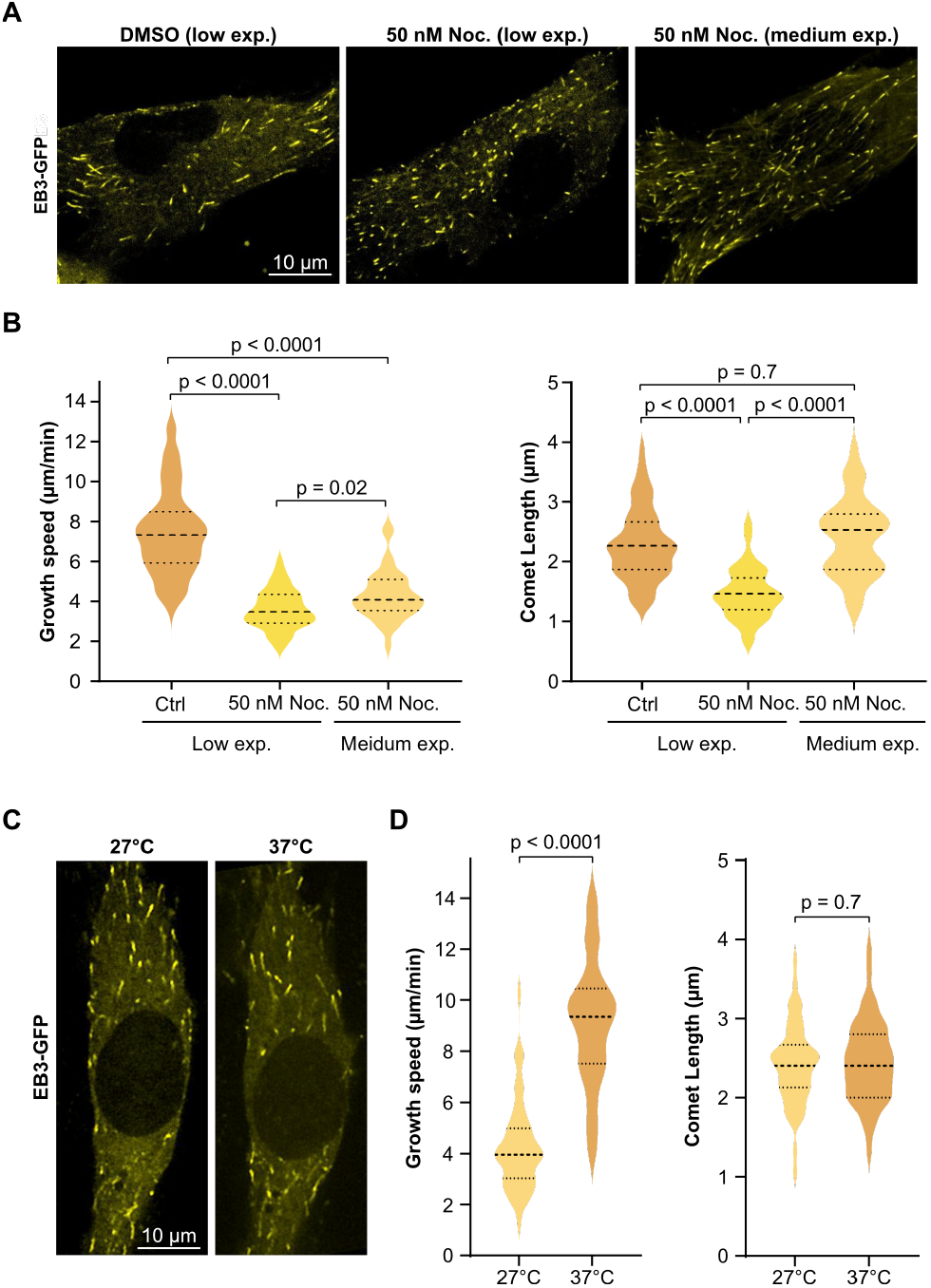
The length of EB3 comets for different microtubule growth velocities. (A) Representative fluorescence images of C2C12 cells transiently transfected with EB3-GFP for control conditions (DMSO low) and in the presence of 50 nM Nocodazole for different EB3-GFP levels (low and medium). (B) Violin plots showing: Microtubule growth speed in the absence and presence of 50 nM Nocodazole for cells with different overexpression levels (left); and corresponding EB3 comets lengths (right). Data collected from: DMSO N=3, 12 cells, 62 comets; 50nMLow N=3, 10 cells, 53 comets; 50nMMedium N=2, 6 cells, 35 comets. Statistical significance was determined using Welch’s t-test for pairwise comparisons (C) Representative fluorescence images of a C2C12 cell transiently transfected with EB3-GFP incubated at *T* = 37^°^C and cooled down to *T* = 27 ^°^C. Violin plots showing: Microtubule growth speed of individual cells analysed first at *T* = 37^°^C and then at *T* = 27^°^C (left); and corresponding EB3 comets lengths (right). Data collected from three independent experiments, with a total of 12 cells analysed and 49 and 44 comets respectively for 27^°^C and 37^°^C. Statistical significance was determined using Welch’s t-test. Scale bars 10 *μ*m.

To further explore the relationship between growth velocity and comet length, we lowered the incubation temperature of low-overexpressing cells from 37^°^C to 27 ^°^C (Fig. 6C,D). In this case, the growth velocity decreased with temperature from 9 *μ*m/min to 4 *μ*m/min, in line with previous observations [33]. However, the EB3 comet length remained constant at around 2.4 *μ*m. Similar results were observed in COS7 cells (SI Appendix, Fig. S7). These findings show that in cells, the EB3 comet length does not scale reliably with microtubule growth velocity. Consequently, the comet length is unlikely to be a measure of the GTP-cap size.

## DROPLET EVAPORATION

*In vitro*, EB3 droplets persist on microtubules throughout the experiment, whereas in cells, remnants vanish within seconds. This discrepancy suggests that a transition from GTP-tubulin to GDP-tubulin along the microtubule is insufficient to trigger EB3 detachment. This observation further supports that the EB3 comet length is not a direct proxy for the GTP-cap size. The behavior of remnants on the microtubule surface aligns with our description of EB3 comets as liquid condensates. Once a droplet or a film has nucleated on a surface, a change of the surface interaction does not result in evaporation of the droplet, because the surface merely accelerates phase separation without altering its intrinsic propensity.

Droplet evaporation can be achieved by introducing two distinct states for the dense phase. In the first state, the material undergoes phase separation as described above; in the second state, phase separation does not occur. The mixture is initially in the phase-separating state and transitions spontaneously to the non-phase-separating state at a rate *k*_*e*_ = 0.07 *s*^−1^. Explicitly, a term −*k*_*e*_*ϕ* is added to our dynamical equation for the phase field, eq. 14. This modification results in droplet evaporation on a timescale similar to that observed in cells (Fig. 7, Movies S7, S8). Our results indicate that an additional reaction process is required to account for the rapid disappearance of remnants *in vivo*.

**FIG. 7.**
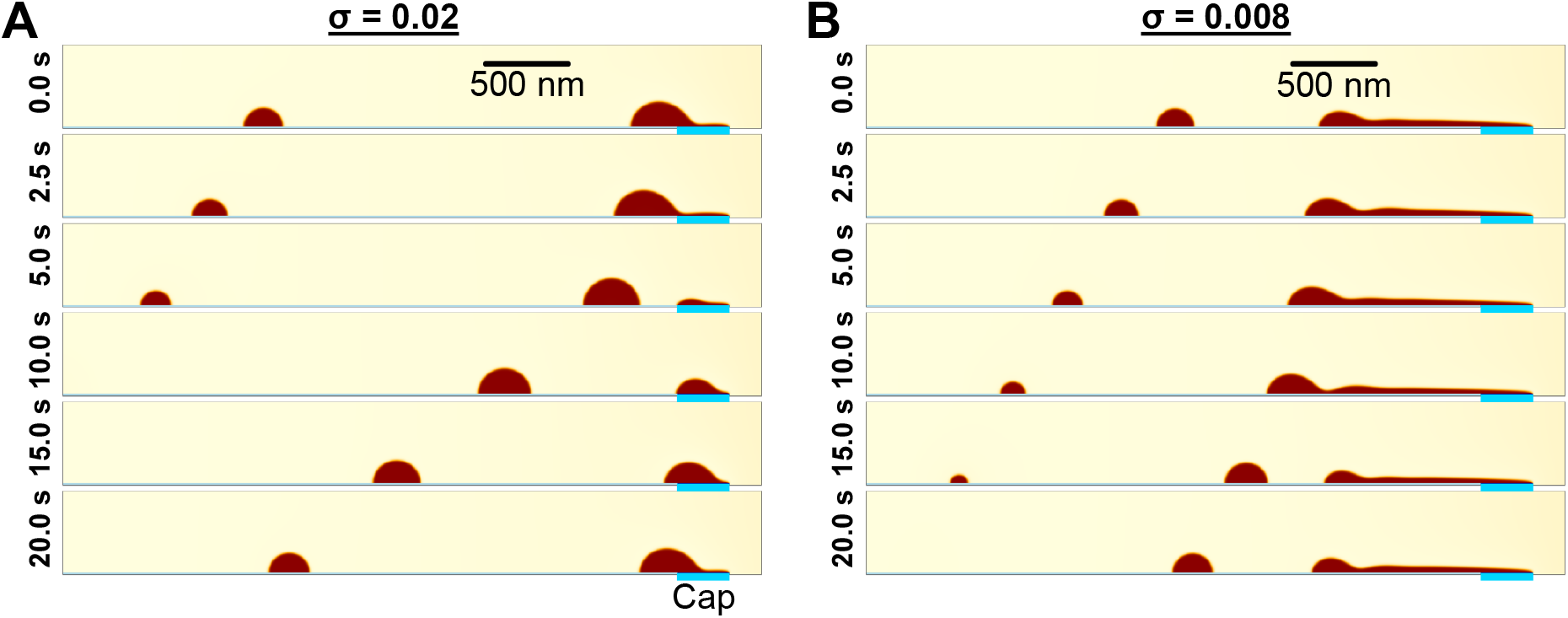
Droplet evaporation at the end of growing cylinders. (A,B) Successive snap-shots of the phase field at the end of a growing cylinder in the droplet (A) and film (B) phases for *v*_MT_ = 8*μm*/*min*. The surface tension is *σ* = 0.02 (A) and *σ* = 0.008 (B). The transition rate is *k*_*e*_ = 0.07*s*^−1^ (A,B). The bulk concentration is *ϕ*_0_ = 0.2 (A,B). Other parameter values as in SI Appendix, Materials and Methods.

## DISCUSSION

In this work, we studied EB3 comet formation on growing microtubules through *in vitro* reconstitution experiments and a physical analysis of condensation on a cylindrical surface. Our results show that EB3 comet formation is compatible with EB3 condensing on the microtubule surface and creating a film that breaks due to the Rayleigh-Plateau instability. We found that the comet length does not depend on the GTP-cap length, but increases with the microtubule growth velocity and the EB3 concentration. Our study highlights the role of non-specific protein-protein interactions in regulating cellular processes.

While our experimental and theoretical results match well, the initial phase of an EB3 film on a microtubule was not accessible experimentally, preventing direct measurements of the time scales *τ*_*m*_ and *τ*_*f*_. Furthermore, we cannot verify directly the relation between the length of the GTP cap and the comet size, because the cap length is not directly accessible. Nonetheless, our results indicate that care should be taken, when the EB3 comet size is taken as a proxy for the cap length.

Future work should focus on understanding how EB3 comet length is regulated in cells. The Rayleigh-Plateau instability might be a key factor, and further unknown factors and proteins might play a role. In contrast to cells, where EB3 remnants rapidly dissolve, EB3 droplets in our *in vitro* experiments persisted. In the context of liquid-liquid phase separation, such dissolution requires a change in internal interactions within the dense phase. In cells, additional cellular factors or a spontaneous conformational change of EB3 as condensates age could alter these interactions.

## MATERIALS AND METHODS

For extended Materials and Methods section see also SI Appendix, Methods.

### Numerical Solution of Dynamic Equations

We numerically solve the Navier-Stokes equation (12) using the Lattice Boltzmann Method (LBM) [32, 34, 35] as described in detail in SI Appendix. The corresponding code can be found at https://github.com/schaerjo/Microtubule-Comets

### Protein purification

EB3 and GFP-EB3 were purified from Escherichia coli BL21 (DE3) (for details, see SI Appendix, Methods). Tubulin was purified as previously described by polymerisation/depolymerisation cycles [36].

## ACKNOWLEDGMENTS

Numeric calculations were performed at the University of Geneva on the “Baobab” HPC cluster. This work was funded by Swiss National Science Foundation Sinergia grant CRSII5 183550, Project grant 310030 212563 and by the DIP of the Canton of Geneva.

## References

[1] C. Duellberg, N. I. Cade, D. Holmes, and T. Surrey, The size of the EB cap determines instantaneous microtubule stabiliTY, eLife 5, e13470 (2016).

[2] A. Akhmanova and M. O. Steinmetz, Microtubule +TIPs at a glance, Journal of Cell Science 123, 3415 (2010).

[3] A. Akhmanova and M. O. Steinmetz, Control of microtubule organization and dynamics: two ends in the limelight, Nat Rev Mol Cell Biol 16, 711 (2015).

[4] G. M. Alushin, G. C. Lander, E. H. Kellogg, R. Zhang, D. Baker, and E. Nogales, High-resolution microtubule structures reveal the structural transitions in αβ-tubulin upon GTP hydrolysis, Cell 157, 1117 (2014).

[5] J. Zhou, A. Wang, Y. Song, N. Liu, J. Wang, Y. Li, X. Liang, G. Li, H. Chu, and H.-W. Wang, Structural insights into the mechanism of GTP initiation of microtubule assembly, Nature Communications 14, 5980 (2023).

[6] D. Roth, B. P. Fitton, N. P. Chmel, N. Wasiluk, and A. Straube, Spatial positioning of EB family proteins at microtubule tips involves distinct nucleotide-dependent binding properties, Journal of Cell Science 132, jcs219550 (2018).

[7] J. Roostalu, C. Thomas, N. I. Cade, S. Kunzelmann, I. A. Taylor, and T. Surrey, The speed of GTP hydrolysis determines GTP cap size and controls microtubule stability, eLife 9, e51992 (2020).

[8] S. P. Maurer, F. J. Fourniol, G. Bohner, C. A. Moores, and T. Surrey, EBs recognize a nucleotide-dependent structural cap at growing microtubule ends, Cell 149, 371 (2012).

[9] R. Zhang, G. M. Alushin, A. Brown, and E. Nogales, Mechanistic origin of microtubule dynamic instability and its modulation by EB proteins, Cell 162, 849 (2015).

[10] S. P. Maurer, N. I. Cade, G. Bohner, N. Gustafsson, E. Boutant, and T. Surrey, EB1 Accelerates Two Conformational Transitions Important for Microtubule Maturation and Dynamics, Current Biology 24, 372 (2014).

[11] A. Urazbaev, A. Serikbaeva, A. Tvorogova, A. Dusenbayev, S. Kauanova, and I. Vorobjev, On the Relationship Between EB-3 Profiles and Microtubules Growth in Cultured Cells, Frontiers in Molecular Biosciences 8, 745089 (2021).

[12] A. Nakano, H. Kato, T. Watanabe, K.-D. Min, S. Yamazaki, Y. Asano, O. Seguchi, S. Higo, Y. Shintani, H. Asanuma, M. Asakura, T. Minamino, K. Kaibuchi, N. Mochizuki, M. Kitakaze, and S. Takashima, AMPK controls the speed of microtubule polymerization and directional cell migration through CLIP-170 phosphorylation, Nat Cell Biol 12, 583 (2010).

[13] H. Bowne-Anderson, M. Zanic, M. Kauer, and J. Howard, Microtubule dynamic instability: A new model with coupled GTP hydrolysis and multistep catastrophe, BioEssays 35, 452 (2013).

[14] F. Perez, G. S. Diamantopoulos, R. Stalder, and T. E. Kreis, CLIP-170 highlights growing microtubule ends in vivo, Cell 96, 517 (1999).

[15] A. Dimitrov, M. Quesnoit, S. Moutel, I. Cantaloube, C. Poüs, and F. Perez, Detection of GTP-Tubulin Conformation in Vivo Reveals a Role for GTP Remnants in Microtubule Rescues, Science 322, 1353 (2008).

[16] C. Aumeier, L. Schaedel, J. Gaillard, K. John, L. Blanchoin, and M. Théry, Self-repair promotes microtubule rescue., Nature Cell Biology 18, 1054 (2016).

[17] H. d. Forges, A. Pilon, I. Cantaloube, A. Pallandre, A.-M. Haghiri-Gosnet, F. Perez, and C. Poüs, Localized Mechanical Stress Promotes Microtubule Rescue, Current Biology 26, 3399 (2016).

[18] V. V. Mustyatsa, A. V. Kostarev, A. V. Tvorogova, F. I. Ataullakhanov, N. B. Gudimchuk, and I. A. Vorobjev, Fine structure and dynamics of EB3 binding zones on microtubules in fibroblast cells, Molecular Biology of the Cell 30, 2105 (2019).

[19] H. Henrie, D. Bakhos-Douaihy, I. Cantaloube, A. Pilon, M. Talantikite, V. Stoppin-Mellet, A. Baillet, C. Poüs, and B. Benoit, Stress-induced phosphorylation of CLIP-170 by JNK promotes microtubule rescue, J Cell Biol 219, e201909093 (2020).

[20] R. Maan, L. Reese, V. A. Volkov, M. R. King, E. O. van der Sluis, N. Andrea, W. H. Evers, A. J. Jakobi, and M. Dogterom, Multivalent interactions facilitate motor-dependent protein accumulation at growing microtubule plus-ends, Nat. Cell Biol. 25, 68 (2023).

[21] S. M. Meier, A.-M. Farcas, A. Kumar, M. Ijavi, R. T. Bill, J. Stelling, E. R. Dufresne, M. O. Steinmetz, and Y. Barral, Multivalency ensures persistence of a +TIP body at specialized microtubule ends, Nat. Cell Biol. 25, 56 (2023).

[22] J. Miesch, R. T. Wimbish, M.-C. Velluz, and C. Aumeier, Phase separation of +TIP networks regulates microtubule dynamics, Proceedings of the National Academy of Sciences of the USA 120, e2301457120 (2023).

[23] X. Song, F. Yang, T. Yang, Y. Wang, M. Ding, L. Li, P. Xu, S. Liu, M. Dai, C. Chi, S. Xiang, C. Xu, D. Li, Z. Wang, L. Li, D. L. Hill, C. Fu, K. Yuan, P. Li, J. Zang, Z. Hou, K. Jiang, Y. Shi, X. Liu, and X. Yao, Phase separation of EB1 guides microtubule plus-end dynamics, Nat. Cell Biol. 25, 79 (2023).

[24] A. A. Hyman, C. A. Weber, and F. Jülicher, Liquid-Liquid Phase Separation in Biology, Annual Review of Cell and Developmental Biology 30, 39 (2014).

[25] X. Zhao, S. Liese, A. Honigmann, F. Jülicher, and C. A. Weber, Theory of Wetting Dynamics with Surface Binding (2024), 2402.10405 [cond-mat, physics:physics].

[26] S. U. Setru, B. Gouveia, R. Alfaro-Aco, J. W. Shaevitz, H. A. Stone, and S. Petry, A hydrodynamic instability drives protein droplet formation on microtubules to nucleate branches, Nat. Phys. 17, 493 (2021).

[27] P. Bieling, S. Kandels-Lewis, I. A. Telley, J. v. Dijk, C. Janke, and T. Surrey, CLIP-170 tracks growing microtubule ends by dynamically recognizing composite EB1/tubulin-binding sites, Journal of Cell Biology 183, 1223 (2008).

[28] P. J. Flory, Thermodynamics of High Polymer Solutions, Journal of Chemical Physics 10, 51 (1942).

[29] M. L. Huggins, Theory of Solutions of High Polymers, Journal of the American Chemical Society 64, 1712 (1942).

[30] D. M. Anderson, G. B. McFadden, and A. A. Wheeler, Diffuse-Interface Methods in Fluid Mechanics, Annual Review of Fluid Mechanics 30, 139 (1998).

[31] P. Yue, J. J. Feng, C. Liu, and J. Shen, A diffuse-interface method for simulating two-phase flows of complex fluids, Journal of Fluid Mechanics 515, 293 (2004).

[32] L. N. Carenza, G. Gonnella, A. Lamura, G. Negro, and A. Tiribocchi, Lattice Boltzmann methods and active fluids, Eur. Phys. J. E 42, 81 (2019).

[33] K. A. Dragestein, W. A. van Cappellen, J. van Haren, G. D. Tsibidis, A. Akhmanova, T. A. Knoch, F. Grosveld, and N. Galjart, Dynamic behavior of GFP-CLIP-170 reveals fast protein turnover on microtubule plus ends, Journal of Cell Biology 180, 729 (2008).

[34] J. G. Zhou, Axisymmetric lattice Boltzmann method, Phys. Rev. E 78, 036701 (2008).

[35] T. Krüger, H. Kusumaatmaja, A. Kuzmin, O. Shardt, G. Silva, and E. M. Viggen, The Lattice Boltzmann Method: Principles and Practice, Graduate Texts in Physics (Springer International Publishing, 2017).

[36] M. Andreu-Carbó, S. Fernandes, M.-C. Velluz, K. Kruse, and C. Aumeier, Motor usage imprints microtubule stability along the shaft, Developmental Cell 57, 5 (2022).

