## Supplementary Information for "Phase Separation and a Hydrodynamic Instability Localize Proteins at Growing Microtubule Ends"

### COMPUTATION OF THE FILM PROFILE

In this section, we compute the profile of a film condensing on a growing cylinder in an infinite medium. To this end, we consider an axisymmetric film condensing on an initially bare cylindrical surface. Neglecting a dependence on the axial coordinate, we compute the radial position $\xi$ of the interface between the condensed and dilute phases as a function of time. With the velocity $v_{\mathrm{MT}}$ of the growing end, we can convert this temporal profile into a spatial profile in the frame of the growing cylinder end.

We assume that the film only grows by diffusion of material from the bulk to the interface. As in the main text, we use the phase field $\phi$ defined in Eq. 1 of the main text to determine the system state. Far from the cylinder, the bulk concentration is $\phi = \phi_0$. For a cylinder of length $l$, mass conservation then implies

$$\partial_t \xi 2\pi\xi l = D\partial_r \phi|_{r=\xi} 2\pi\xi l \tag{1}$$

and thus

$$\tag{2}$$

$$\partial_t \xi = D\partial_r^2 \phi|_{r=\xi}. \tag{3}$$

Outside the condensed phase, the dynamics of the phase field $\phi$ is given by

$$\partial_t \phi = D\left(\partial_r^2 \phi + \frac{1}{r}\partial_r \phi\right). \tag{4}$$

Here, $D = k_B \bar{T} M$ is the diffusion constant, wheer $k_B$ is the Boltzmann constant and $\bar{T}$ absolute temperature. In the following we use units with $k_B \bar{T} = 1$. Material reaching the interface is immediately absorbed, which means that $\phi\,(r = \xi) = 0$. Following Ref. [1], we rescale the variables as $X = \frac{r}{r_i} - 1$, $\delta = \frac{\xi}{r_i} - 1$ and $T = \frac{tD}{r_i^2}$, where $r_i$ is the radius of the


* co-first author

†

‡

microtubule. The system of equations to solve then reads

$$\partial_T\phi = \partial_X^2\phi + \frac{\partial_X\phi}{1+X} \tag{5}$$

$$\phi\left(X, T=0\right) = \phi_0 \tag{6}$$

$$\phi\left(X=\delta\left(T\right), T\right) = 0 \tag{7}$$

$$\phi\left(X=\rightarrow\infty, T\right) = \phi_0 \tag{8}$$

$$\delta\left(0\right) = 0 \tag{9}$$

$$\partial_T\delta = \partial_X\phi|_{X=\delta}. \tag{10}$$

We now introduce the similarity transformation $\eta = X/\delta$ and demand $\phi = \phi\left(\eta\right)$. Equation 5, with $\partial_\eta\phi = \phi'$, becomes

$$-\eta\frac{1}{\delta}\frac{d\delta}{dT}\phi' = \frac{\phi''}{\delta^2} + \frac{\phi'}{\delta\left(1+\eta\delta\right)}. \tag{11}$$

For small $T$ and near the interface, we have $\eta = \mathcal{O}\left(1\right)$ and $\delta \ll 1$. Thus, we can neglect $\eta\delta$ when compared to 1, which transforms Eq. 11 into

$$\phi'' = -\phi'\left(\eta\delta\frac{d\delta}{dT} + \delta\right). \tag{12}$$

We also assume that $\mathcal{O}\left(\frac{d\delta}{dT}\right) > \mathcal{O}\left(\delta\right)$, thus we neglect the isolated $\delta$, which transforms Eq. 12 into

$$\phi'' = -\eta\phi'\left(\eta\delta\frac{d\delta}{dT}\right). \tag{13}$$

To satisfy $\phi = \phi\left(\eta\right)$, we need $\delta\frac{d\delta}{dT} = b$ with $b$ a constant. By using Eq. 9 the solution is

$$\delta = \sqrt{2bT}. \tag{14}$$

We will now solve Eq. 13. First, we rewrite the boundary conditions Eqs. 6 and 8 as $\phi\left(\eta\rightarrow\infty\right) = \phi_0$ and Eq. 7 as $\phi\left(\eta=1\right) = 0$. The solution of Eq. 13 then is

$$\phi\left(\eta\right) = \sqrt{\frac{\pi}{2b}}A_1\mathrm{erf}\left(\eta\sqrt{b/2}\right) + A_2. \tag{15}$$

where $A_1$ and $A_2$ are integration constants and $\mathrm{erf}\left(x\right)$ is the error function. The expressions of these constants are obtained by using the boundary conditions and we get

$$\phi\left(\eta\right) = \phi_0 - \phi_0\frac{1-\mathrm{erf}\left(\eta\sqrt{b/2}\right)}{1-\mathrm{erf}\left(\sqrt{b/2}\right)}. \tag{16}$$

The expression of $b$ is obtained self-consistently by using $b = \delta \frac{d\delta}{dT} = \phi'|_{\eta=1}$ :

$$\frac{\sqrt{\pi}}{\phi_0}\sqrt{b/2} = \frac{e^{-b/2}}{1 - \operatorname{erf}\left(\sqrt{b/2}\right)}. \tag{17}$$

Finally, transforming Eq. 14 back into the original variables, the time evolution of the interface position $\xi$ is given by

$$\xi\,(t) = r_i + \sqrt{2bDt} \tag{18}$$

with $b$ given by Eq. 17. To determine the profile of the film near the tip, we use Eq. 18 with $\zeta = v_{MT}t$, where $\zeta$ is the distance from the tip along the cylinder axis. For the parameters chosen in the main text, the analytical result matches well the numerical interface profile (Fig. 2A).

### MATERIALS AND METHODS

#### Numerical Solution of the Dynamic Equations

We numerically solve the Navier-Stokes equation 12 of the main text using the Lattice Boltzmann Method (LBM) [2–4]. We use cylindrical coordinates $(r, \theta, z)$. Assuming axial symmetry, we solve the dynamic equations on a square lattice in the $(r, z)$-plane. In the LBM, the fluid is described in terms of the partial distribution functions $f_i$ defined on the lattice sites $\mathbf{r}_n$ and for the discrete velocities $\boldsymbol{\xi}_i = c\mathbf{e}_i$, where $c = \Delta x/\Delta t$ with $\Delta x$ being the lattice spacing and $\Delta t$ the time step. We choose a set of 9 velocities (D2Q9) with $\mathbf{e}_0 = (0, 0)$, $\mathbf{e}_{1,3} = (\pm 1, 0)$, $\mathbf{e}_{2,4} = (0, \pm 1)$, and $\mathbf{e}_{5-8} = (\pm 1, \pm 1)$. From the distribution functions $f_i$, the fluid density $\rho$ and velocity $\mathbf{u}$ are obtained via

$$\rho = \sum_i f_i \tag{19}$$

$$\rho\mathbf{u} = \sum_i \boldsymbol{\xi}_i f_i + \frac{\Delta t}{2}\mathbf{F}_s, \tag{20}$$

where $\mathbf{F}_s$ is the interfacial force.

The distribution functions evolve according to

$$f_i\left(\mathbf{r}_n + \boldsymbol{\xi}_i \Delta t, t + \Delta t\right) - f_i\left(\mathbf{r}_n, t\right) = \left\{\mathcal{C}_i + S_i + \mathcal{S}_i\right\}\Delta t \tag{21}$$

representing streaming followed by a collision step. The collision step has several contributions. First, assuming a single relaxation time $\tau$

$$\mathcal{C}_i = -\frac{1}{\tau}\left(f_i\left(\mathbf{r}_n, t\right) - f_i^{\mathrm{eq}}\left(\mathbf{r}_n, t\right)\right), \tag{22}$$

where the local equilibrium distribution function $f_i^{\mathrm{eq}}$ is

$$f_i^{\mathrm{eq}}\left(\mathbf{r}_n, t\right) = \omega_i \rho \left(1 + \frac{\mathbf{u}\cdot\boldsymbol{\xi}_i}{c_s^2} + \frac{\left(\mathbf{u}\cdot\boldsymbol{\xi}_i\right)^2}{2c_s^4} - \frac{\mathbf{u}\cdot\mathbf{u}}{2c_s^2}\right). \tag{23}$$

Here, $c_s$ is the speed of sound with $c_s^2 = (1/3)\Delta x^2/\Delta t^2$. Second, the interfacial force $\mathbf{F}_s$ leads to the term

$$S_i = \left(1 - \frac{\Delta t}{2\tau}\right)\omega_i\left(\frac{\xi_{i,\alpha}}{c_s^2} + \frac{\left(\xi_{i,\alpha}\xi_{i,\beta} - c_s^2\delta_{\alpha\beta}\right)u_\beta}{c_s^4}\right)F_\alpha, \tag{24}$$

where $F_\alpha$ are the components of the force density and summation over greek indices is assumed. Finally, there are additional source terms resulting from using cylindrical coordinates. Explicitly,

$$\mathcal{S}_i = \Theta + \frac{1}{6c^2}\xi_{i,\alpha}F_\alpha^{\mathrm{axi}} \tag{25}$$

with

$$\Theta = -\frac{\rho u_r}{9r} \tag{26}$$

and

$$F_\alpha^{\mathrm{axi}} = -\frac{\rho u_\alpha u_r}{r} + \frac{\nu}{r}\partial_r u_\alpha - \frac{\nu u_\beta}{r^2}\delta_{\beta r}. \tag{27}$$

The derivatives of the velocity $\mathbf{u}$ in Eq. 27 are computed via [2]

$$\partial_r u_x = -\frac{3}{2\rho\tau c^2\Delta t}\sum_i \xi_{i,x}\xi_{i,r}\left(f_i - f_i^{\mathrm{eq}}\right) - \partial_x u_r \tag{28}$$

$$\partial_r u_r = -\frac{3}{2\rho\tau c^2\Delta t}\sum_i \xi_{i,r}\xi_{i,r}\left(f_i - f_i^{\mathrm{eq}}\right) - \frac{u_r}{2r} \tag{29}$$

to improve stability and reduce the computational cost.

The Cahn-Hillard equation 2 in the main text for the phase field $\phi$ is solved numerically by a finite difference upwind scheme.

**Simulation parameters**

In the Lattice-Boltzmann Method, we use $\rho = 1$, $\Delta t = 1$ and $\Delta x = 1$. To convert them into physical units, we choose a length scale of 4 nm and a time scale of 10 s $10^{-4}$. For solving the diffusion equation, we use $\Delta t^{\text{diff}} = 0.01$ or 0.1 depending on the value of $M\sigma$.

The parameters of the simulations are the mobility $M$, the surface tension $\sigma$, the interface width $\epsilon$, the microtubule growth velocity $v_{MT}$, the GTP-cap length $\ell_{\text{GTP}}$, the initial bulk concentration $\phi_0$, the kinematic viscosity $\nu$ and the interaction strength $h$. For all simulations, we chose $\epsilon = 2$ and $h/\kappa = 0.4$, such that a layer of the dense phase is present near the attractive surface.

Unless stated otherwise, we used the following parameter values. We chose $M = 31.25$ and $\sigma = 0.001$. This value of $\sigma$ and the variations considered in the simulations correspond to a physical value of $\sigma^{phy} \sim 10^{-3} - 10^{-5}\, mN/m$, which is similar to values measured for biomolecular condensates [5]. Furthermore, $v_{MT} = 4.17 \cdot 10^{-4}$ which corresponds to $v_{MT} = 1\, \mu m/min$ and is similar to the growth speed observed *in vitro* (cite). Also, $\ell_{\text{cap}} = 80$, which corresponds to $360\, nm$ and $\phi_0 = 0.05$. These values were chosen to ensure consistent nucleation at the tip and to avoid discretization artifacts. In the Lattice Boltzmann Method, the kinematic viscosity $\nu$ is related to the relaxation time $\tau$ via $\nu = (\tau - 0.5)\, c_s^{-2}$. For numerical stability and precision, one should use $\tau \approx 1$ as a general rule. We took $\tau = 0.9$. This value leads to a Reynolds number $Re = uL/\nu \approx 10^{-2}$, where $u \approx 10^{-4}$ is the maximal fluid velocity achieved in our simulations and $L \approx 100$ is the radial domain size. As a consequence, inertial effects are negligible.

**Cell culture and treatments**

COS-7 and C2C12 cells were cultured in high glucose Dulbecco's Modified Eagle's Medium (DMEM, ThermoFisher) supplemented with 10 % Fetal Bovine Serum (FBS, ThermoFisher, 10270106) and 1 % penicillin-streptomycin (Gibco, 15140122) at 37$°C$ with 5 % CO2. Cells were transfected with 0.5 $\mu g$ and 1 $\mu g$ EB3-GFP respectively using the jetOPTIMUS transfection reagent (Polyplus) according to the manufacturer's instructions. The transfection medium was changed before imaging, 15h after transfection. To affect microtubule polymerization speed, cells were treated with 50 $nM$ nocodazole (Sigma, M1404;

diluted in culture medium) for 1 hour before imaging.

#### Cloning

EB3-GFP (mammalian expression) was generated by excising GFP from an EB3-mCherry (Addgene plasmid 55037) vector using AgeI/BsrG1 restriction sites and replacing it with GFP containing AgeI/BsrG1 overhangs generated by PCR. The pet28a-6XHis-EB3 and pet28a-GFP-EB3-6Xhis were used in a previous publication. The pet28a-6XHis-EB3 were kind gifts of Natacha Olieric.

#### Imaging

*Confocal Microscopy*

For in vitro microtubule dynamics experiments, imaging was performed on an Axio Observer Inverted TIRF microscope (Zeiss, 3i) equipped with a Prime 95B BSI (Photometrics) using a 100X objective (Zeiss, Plan-Apochromat 100X/1.46 oil DIC (UV) VIS-IR). SlideBook 6 X 64 software (version 6.0.22) was employed to record time-lapse imaging. To keep microtubules at $37°C$, the stage was provided with a Chamlide Live Cell Instrument incubator ($37°C$ for in vitro experiments, supplemented with 5 % CO2 for live cell experiments). For In-cell studies, imaging was performed with a 3i Marianas spinning disk confocal setup based on a Zeiss Z1 stand, a 100× PLAN APO NA 1.45 TIRF objective, and a Yokogawa X1 spinning disk head followed by a 1.2× magnification lens, and an Evolve EMCCD camera (Photometrics). A stage-top incubator (Okolab) was used for cells to keep them at $37°C$ and to decrease incubation temperature to $27°C$ .

*Negative stain transmission electron microscopy*

For negative stain transmission electron microscopy (negative stain EM), stabilized microtubule seeds were diluted 1:60 in BRB1X buffer supplemented with 60 mM KCl. EB3 was added to the diluted sample to a final concentration of 5 $\mu$M. A 4 $\mu$L aliquot of the sample was applied directly onto glow-discharged copper EM grids (Electron Microscopy Sciences). After a one-minute incubation, the excess sample was blotted off, and the grids were washed twice with 10 $\mu$L drops of 2% uranyl acetate (Electron Microscopy Sciences), each time followed by immediate blotting. The grids were then stained with a 10 $\mu$L drop of 2% uranyl acetate for one minute, blotted, and air-dried for two minutes. Prepared grids were stored in EM grid boxes at room temperature until imaging. Imaging was performed using a Talos L120C transmission electron microscope.

*Cryo-electron microscopy*

For cryo-ET experiments, seeds were diluted 1:30 in BRB1X buffer supplemented with 60 $mM$ KCl and 5 $\mu M$ EB3. After sample pipetting and 30 s incubation onto the grid, it was backblotted at room temperature with 90 % humidity before plunging into liquid ethane using an EM GP2 automatic plunge freezer (Leica). The grid was stored in liquid nitrogen until image acquisition. Cryo-ET tilt series data was acquired on a Talos Arctica electron microscope operated at 200kV, equipped with a Falcon 4i Direct Electron Detector and Selectris X energy filter (all ThermoFisher Scientific). Data was recorded using Tomography 5.10 software (ThermoFisher Scientific). The pixel size was 2.5 Angstrom, and the tiles were $\pm$ 60$^{\circ}C$ with 3 degrees between each tilt.

**Tubulin purification from bovine brain and labelling**

Tubulin was purified as described previously in [6] from fresh bovine brains after two depolymerization/polymerization cycles. Tubulin was labeled with biotin and ATTO-565 as described in [6].

**Protein purification**

Purification of EB3 and GFP-EB3 were performed in E.coli B121 cells as previously published in [7].

**Coverslip treatment and Flow chamber preparation**

For in vitro microtubule studies, slides and coverslips were cleaned by 1M NaOH and sonicated for 30 min. They were then washed in a beaker of distilled water. They were then sonicated in 96% ethanol for 30 minutes and again washed in distilled water. After careful drying with an air gun, they were plasma-cleaned. They were then incubated for 48 with tri-ethoxy-silane-PEG (Creative PEGWorks) or a 1:5 mix of tri-ethoxy-silane-PEG-biotin: tri-ethoxy-silane-PEG (final concentration 1 mg/ml) in 96 % ethanol and 0.02 % HCl, with gentle agitation at room temperature. They were washed in 96 % ethanol, washed in distilled water, dried with the air gun, and kept in the fridge for up to one week. Flow chambers were prepared by adding double-sided on the slides and tapping the coverslips.

**Microtubule dynamics assays in vitro**

Microtubule seeds were prepared using 10 $\mu M$ tubulin (30% biotinylated and 70% ATTO-647) with 0.5 $mM$ GMPCPP in BRB80 buffer. After 30 min incubation in the 37°$C$ bath 1 $\mu M$ taxol was added and the solution was incubated for 30 more minutes. The solution was then centrifuged at 14 000 g for 15 min at room temperature to pellet the seeds. They were then resuspended in BRB80 supplemented with 0.5 $mM$ GMPCPP and 1$\mu M$ taxol. They were then aliquoted and kept in liquid nitrogen.

For microtubule in vitro assay, the flow chamber was first injected with 50 $\mu g/mL$ neutravidin (Thermofisher) in BRB80, then washed with BRB80 and incubated with seeds. The chamber was then injected three times with BRB/BSA. Then together with anti-bleaching buffer (10 $mM$ DTT, 0.3 $mg/mL$ glucose, 0.1 $mg/mL$ glucose oxidase, 0.02 $mg/mL$ catalase, 0.125 % methylcellulose (1500 cP, Sigma), 1 $mM$ GTP), 10 $\mu M$ tubulin ATTO-565 labelled-tubulin (1:5 ratio labeled to unlabeled) and the appropriate salt, protein and polyethylene glycol (PEG) 4000.

For the experiments with only seeds and the "low salt" experiments, 5, 10, and 20 $\mu M$ EB3 were used with 60 $mM$ KCl. In this case, 90% of unlabeled EB3 was used with 10% GFP-EB3. For the "high salt" experiments, 700 $nM$ GFP-EB3 was used together with 2% PEG, 60 $mM$ KCl, and 85 $mM$ Kacetate. It has to be noted that antibleaching buffer and the presence of 100% labelled GFP drastically diminish the phase separation behavior of EB3 (Supplementary Figure X). Due to this in the "tip-tracking" condition we used PEG as a crowding agent. A higher salt concentration is used to induce tip tracking and to inhibit electrostatic interactions with the rest of the microtubule.

#### Microtubule dynamics

For in-cell and in-vitro microtubule dynamics measurements, images were taken every 3s. Comets were tracked using the Line tool in ImageJ, and kymographs were built using the KymographBuilder plugin. Lines were drawn on the kymograph to get the slope, and a custom-written macro was used to determine microtubule growth speed.

#### Comet length analysis

For comet length analysis in cells, we define low-, and medium-, overexpressing cells as cells that exhibit mean fluorescence intensity within a range of respectively 2000 to 3000 A.U., and 3000-9000 A.U.. Cells with higher fluorescent intensity were excluded from the analysis. Comets for the analysis were chosen so that they would not overlap with each other and would be visible for at least 15 frames. For a given comet, the comet length was measured by taking the longest value of the timelapse to minimize errors due to comets moving out of the focus plane. The background from the microscope imaging was of 1000 A.U.. To measure the length of each comet, we perform a linescan along the comet axis. We determine the local background by adding the microscope background to the minimum value of the linescan. This value is taken as a threshold to determine the comet region and thus its length.

#### Fluorescence recovery after photobleaching

Fluorescence recovery after photobleaching (FRAP) experiments in cells were performed with a 656 $nm$ laser at 100% laser intensity to be sure to get rid of all the fluorescence. The analysis was done using the formula

$$F_{norm}(t) = \frac{F_{ROI}(t) - F_{bck}}{F_{ctrl}(t) - F_{bck}} \frac{F_{ctrl}(i) - F_{bck}}{F_{ROI}(i) - F_{bck}} \tag{30}$$

where $F_{ROI}(t)$ and $F_{ctrl}(t)$ are respectively the ROI and the control fluorescence intensity before the FRAP, $F_{bck}$ the background fluorescence and $F_{ROI}(i)$ and $F_{ctrl}(i)$ are respectively the ROI of the unbleached part of the condensate at one timepoint (i)[8]

**Statistical analysis**

For statistical analysis, GraphPad Prism software 10 was used.

**Movie S1.** EB3 comet with remnant formation and evaporation in C2C12 cells untreated at $37°C$.

**Movie S2.** EB3 comet with remnant formation and evaporation in C2C12 cells untreated at $37°C$.

**Movie S3.** Movie corresponding to the data presented in Figure 2A. The attractive cap is represented in black. Dashed line: saturated height $\xi_\infty$. Scale bar: $500\,nm$.

**Movie S4.** Movie corresponding to the data presented in Figure 2B. The attractive cap is represented in black. Scale bar: $500\,nm$.

**Movie S5.** Movie corresponding to the data presented in Figure 4A. The attractive cap is represented in black. Scale bar: $500\,nm$.

**Movie S6.** Movie corresponding to the data presented in Figure 5A. The attractive cap is represented in black. Scale bar: $500\,nm$.

**Movie S7.** Movie corresponding to the data presented in Figure 7A. The attractive cap is represented in black. Scale bar: $500\,nm$.

**Movie S8.** Movie corresponding to the data presented in Figure 7B. The attractive cap is represented in black. Scale bar: $500\,nm$.

---


[1] S. U. Setru, B. Gouveia, R. Alfaro-Aco, J. W. Shaevitz, H. A. Stone, and S. Petry, A hydrodynamic instability drives protein droplet formation on microtubules to nucleate branches, Nat. Phys. **17**, 493 (2021).

[2] J. G. Zhou, Axisymmetric lattice Boltzmann method, Phys. Rev. E **78**, 036701 (2008).

[3] T. Krüger, H. Kusumaatmaja, A. Kuzmin, O. Shardt, G. Silva, and E. M. Viggen, *The Lattice Boltzmann Method: Principles and Practice*, Graduate Texts in Physics (Springer International Publishing, 2017).

[4] L. N. Carenza, G. Gonnella, A. Lamura, G. Negro, and A. Tiribocchi, Lattice Boltzmann methods and active fluids, Eur. Phys. J. E **42**, 81 (2019).

[5] H. Wang, F. M. Kelley, D. Milovanovic, B. S. Schuster, and Z. Shi, Surface tension and viscosity of protein condensates quantified by micropipette aspiration, Biophys. Rep. **1**, 100011 (2021).

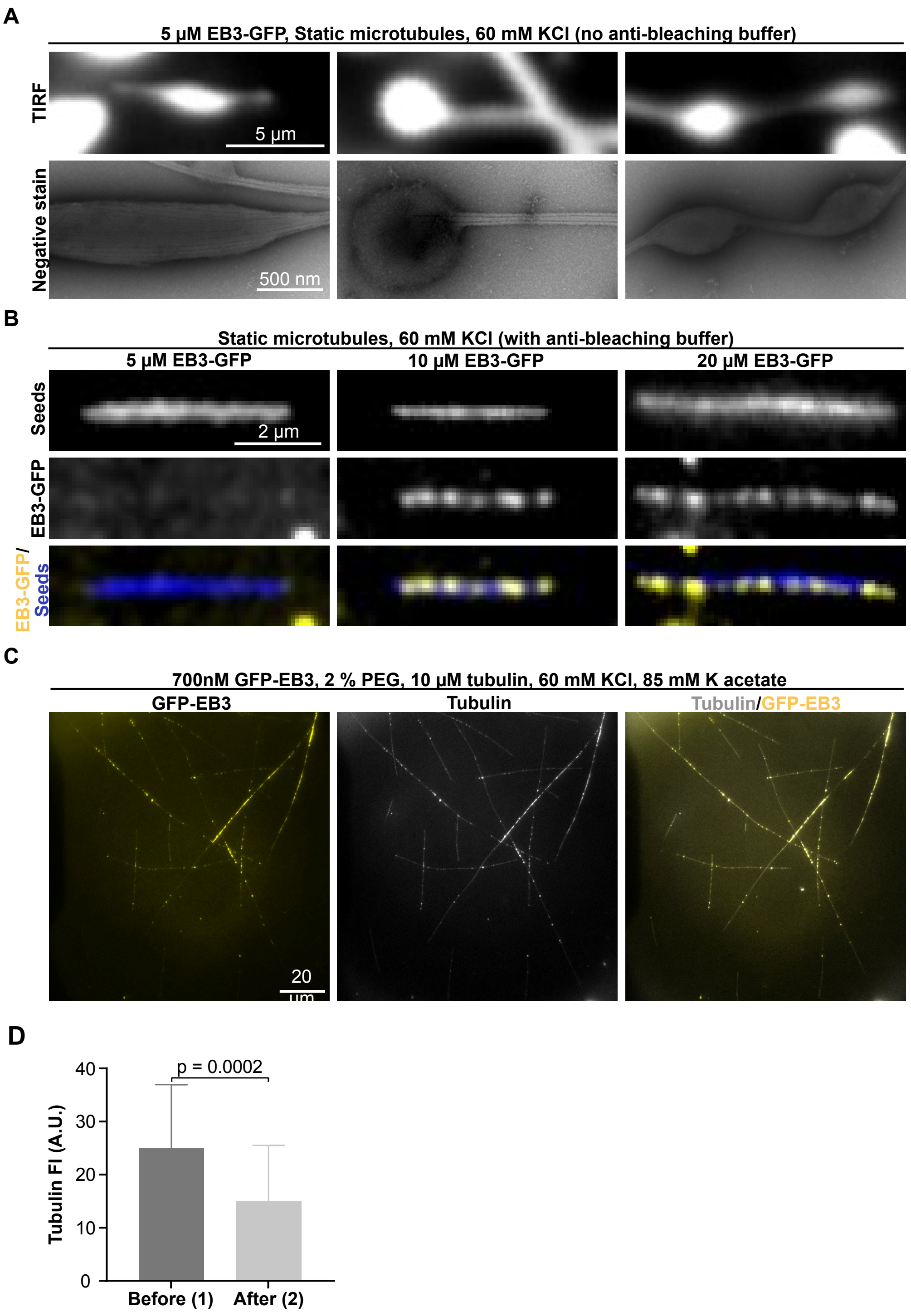


FIG. S1. Effect of concentration and anti-bleaching buffer on EB3 condensation. (A) Representative TIRF (top) and negative staining (bottom) images of EB3 at 5 $\mu$M without anti-bleaching buffer on static microtubules. EB3 wets the microtubule and forms condensates. (B) Representative TIRF images of EB3 5 $\mu$M, 10 $\mu$M, and 20 $\mu$M on static microtubules with anti-bleaching buffer. (C) Representative TIRF images of growing microtubules under high salt concentration (similar to Fig. 1) showing phase separation only occurs on microtubules and not in the bulk. (D) Tubulin fluorescence at the growing microtubule end just before and after remnant fission.

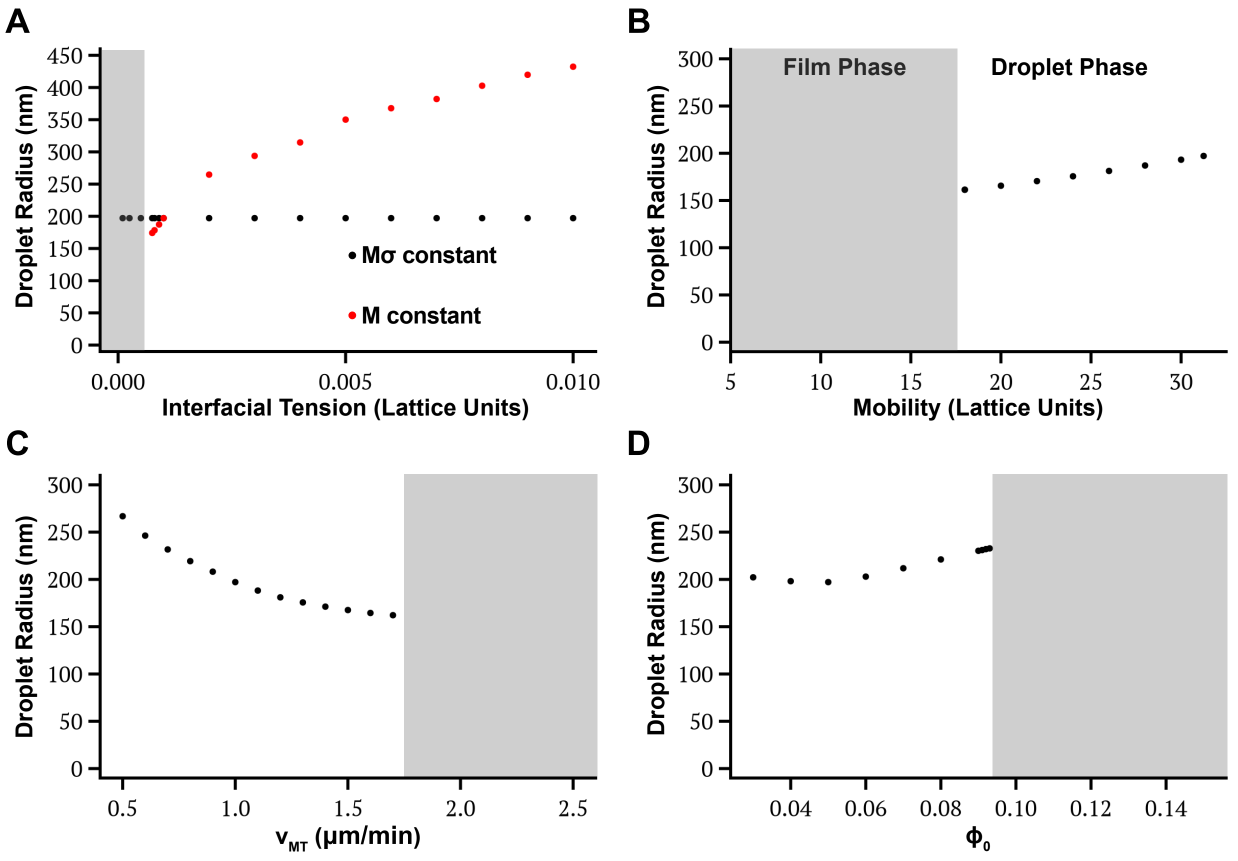


FIG. S2. Average droplet radius on growing cylinders (theory) as a function of surface tension $\sigma$ (A), mobility $M$ (B), cylinder growth velocity $v_{MT}$ (C) and bulk phase field value $\phi_0$ (D). In (A), the surface tension is varied while keeping $M\sigma$ (black) or $M$ (red) constant.The grey shaded areas indicate existence of the film phase. Other parameter values are given in Materials and Methods, Simulation parameters.


[6] M. Andreu-Carbó, S. Fernandes, M.-C. Velluz, K. Kruse, and C. Aumeier, Motor usage imprints microtubule stability along the shaft, Developmental Cell **57**, 5 (2022).

[7] J. Miesch, R. T. Wimbish, M.-C. Velluz, and C. Aumeier, Phase separation of +TIP networks regulates microtubule dynamics, Proceedings of the National Academy of Sciences of the USA **120**, e2301457120 (2023).

[8] C. A. Day, L. J. Kraft, M. Kang, and A. K. Kenworthy, Analysis of protein and lipid dynamics using confocal Fluorescence Recovery After Photobleaching (FRAP), Current Protocols in Cytometry **62**, 2.19.1 (2012).

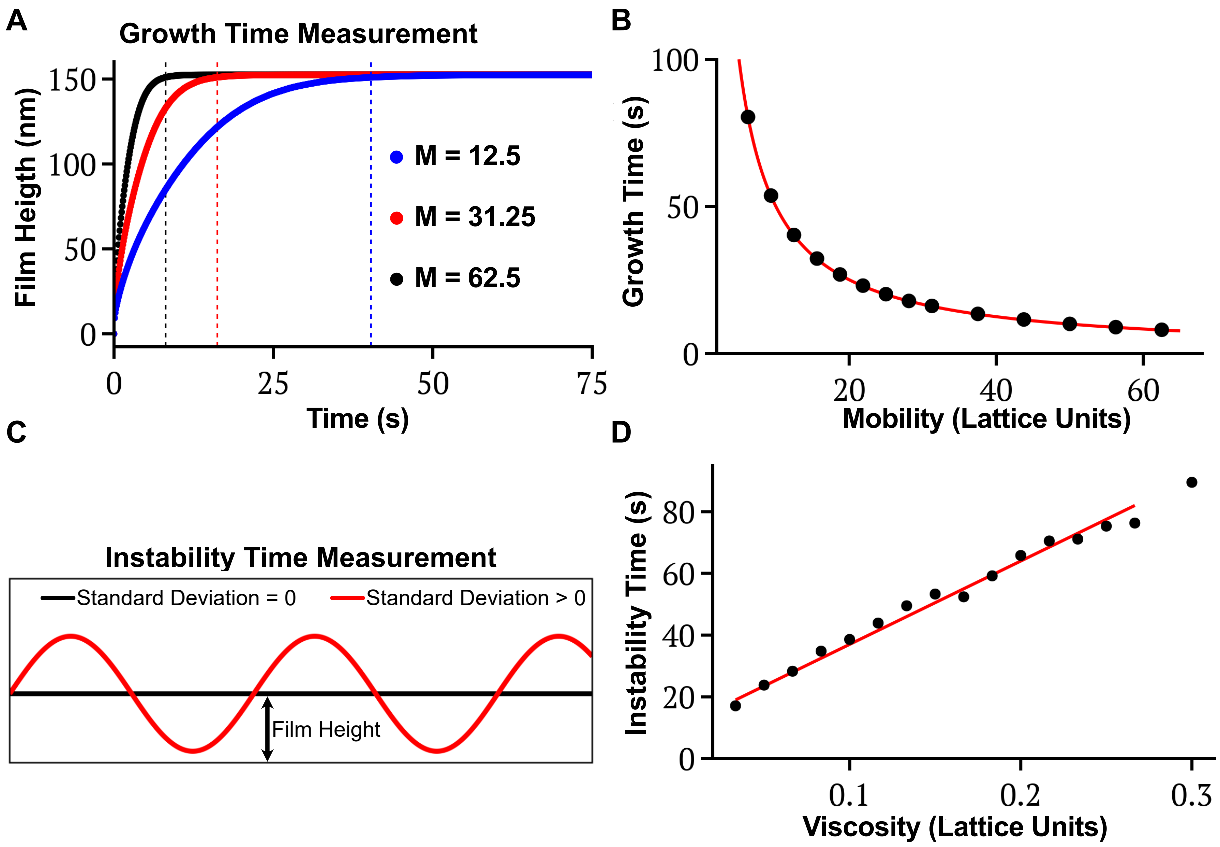


FIG. S3. Droplet formation on static microtubules. (A) Film height as a function of time for three different mobilities. The vertical dotted lines are the time points at which the average film height reaches 95% of its final value. These points are used to measure the growth time $\tau_f$. (B) Growth time of the film $\tau_f$ as a function of the mobility $M$. (C) Scheme representing the evolution of the interface during the simulations to measure the instability time $\tau_m$. The black line is the initial state, a cylindrical film with a constant radius and thus no standard deviation of the interface height. The red line represent the interface after the Rayleigh-Plateau instability has developed and thus a finite standard deviation. The time $\tau_m$ is defined as the time when the standard deviation has reached 5% of the initial film height. (D) The scale $\tau_m$ as a function of the viscosity $\nu$. The red lines are guides to the eye of the form $a/M$ (B) and $a\nu + b$ (D), where $a$ and $b$ are constants. The simulation parameters are the ones described in the Materials and Methods section.

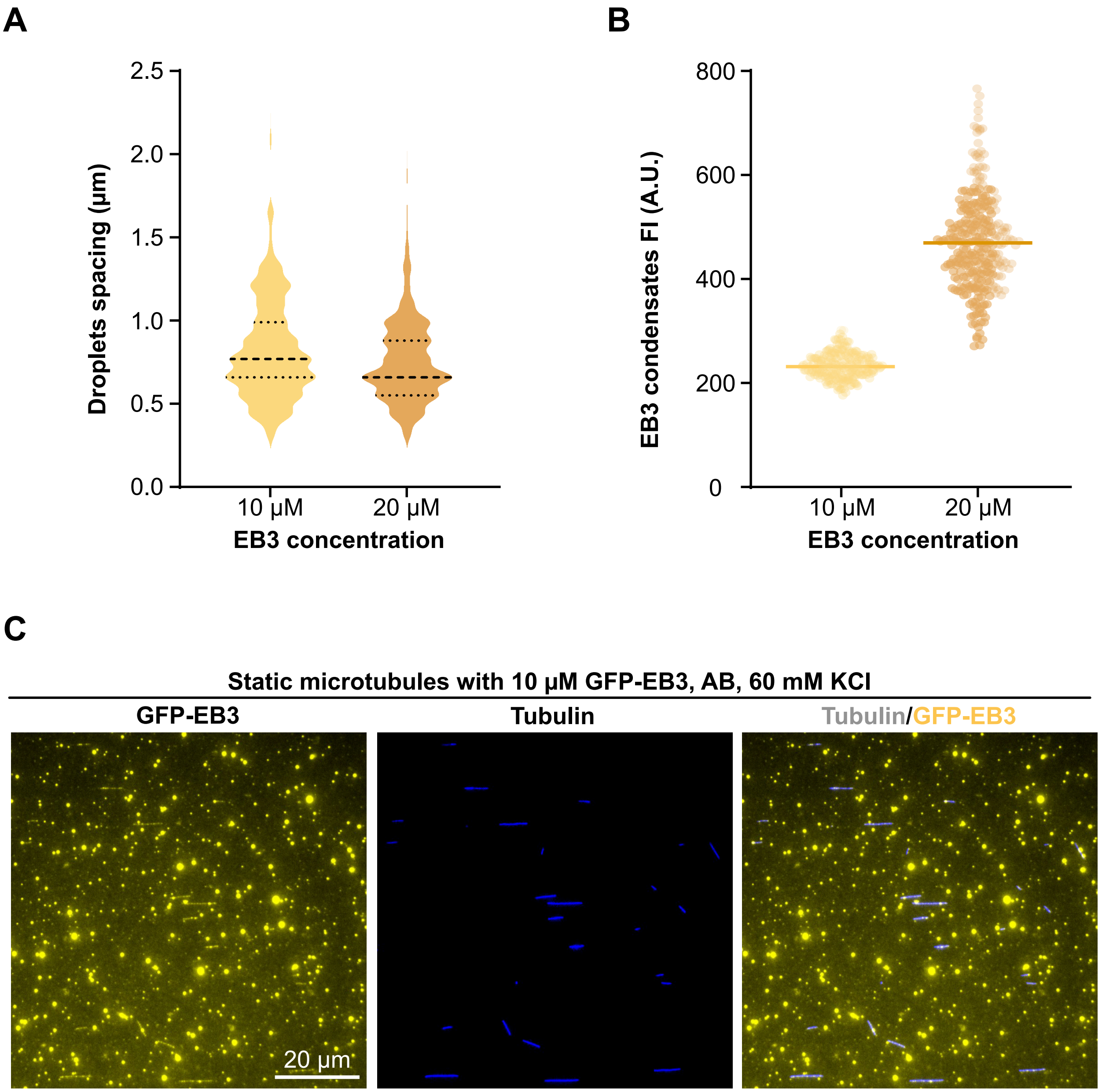


FIG. S4. EB3 condensation on static microtubules and in the bulk. (A) Inter-EB3-droplet distance for two concentrations of EB3. (B) Fluorescence intensity of individual EB3 droplets along microtubules for two concentrations. (C) Representative TIRF images of static microtubules in presence of 10 $\mu M$ GFP-EB3 showing EB3 condensation on microtubules and in the bulk.

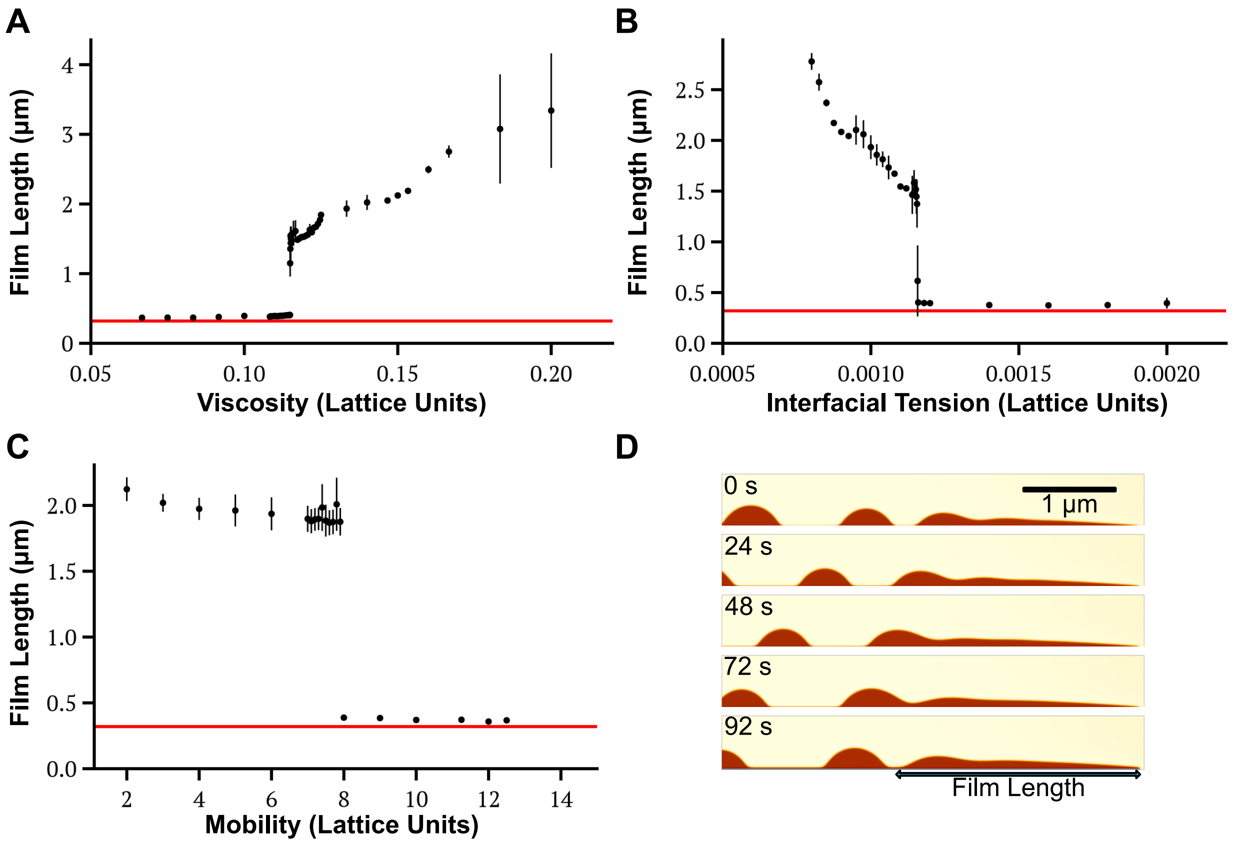


FIG. S5. Transition between droplet and film phases as function of the model parameters. (A-C) Average film length as a function of the viscosity $\nu$ (A), the interfacial tension $\sigma$ (B) and the mobility $M$ (C). The red lines represent $\ell_{\text{cap}} = 320\,\text{nm}$. (D) Representative snapshots from a simulation showing condensation on a growing cylinder with flows induced by interfacial gradients for a uniformly attractive cylinder. The simulation parameters are the ones described in the Materials and Methods section except for $M = 6.25$.

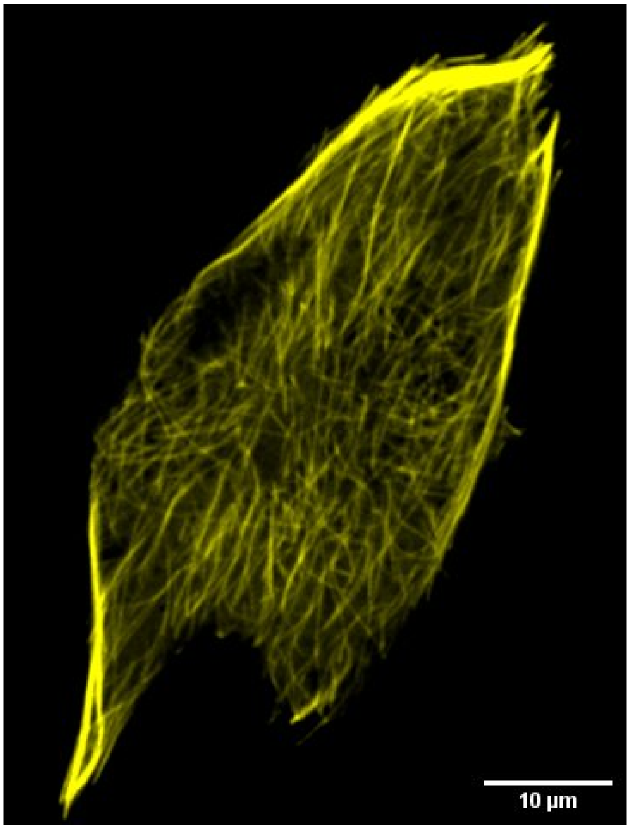


FIG. S6. Representative fluorescence image of C2C12 cell transiently transfected with EB3-GFP showing high level of overexpression. At high overexpression levels EB3 covers the complete microtubule.

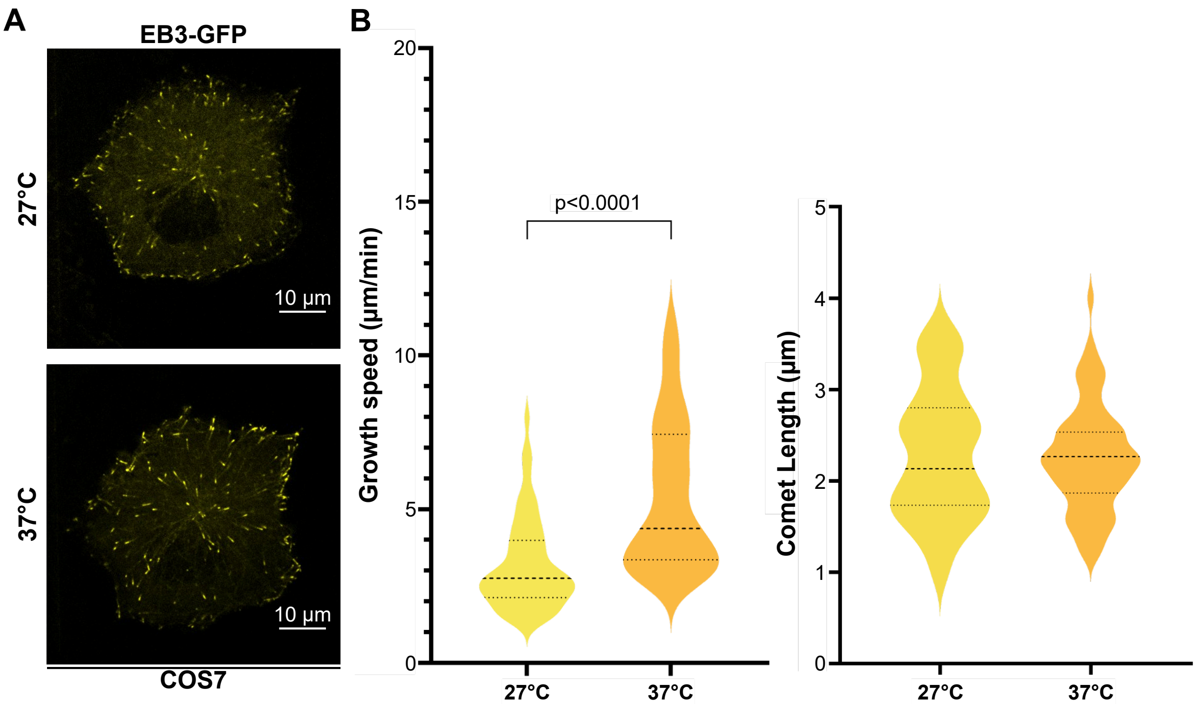


FIG. S7. The length of EB3 comets does not scale with the microtubule growth velocity. (A) Representative fluorescence images of a COS7 cell transiently transfected with EB3-GFP incubated at $T = 37\,°\mathrm{C}$ and cooled down to $T = 27\,°\mathrm{C}$ . (B) Violin plots showing: Microtubule growth speed of individual cells analysed first at $T = 37\,°\mathrm{C}$ and then at $T = 27\,°\mathrm{C}$ (left); and corresponding EB3 comets lengths (right). Data collected from three independent experiments, with a total of 52 comets analysed from 9 cells for 37 °C condition and 57 comets from 7 cells for 27 °C. Statistical significance was determined using Welch's t-test. Scale bars $10\,\mu$m.
